# Enriching and aggregating purple non-sulfur bacteria in an anaerobic sequencing-batch photobioreactor for nutrient capture from wastewater

**DOI:** 10.1101/2020.01.08.899062

**Authors:** Marta Cerruti, Berber Stevens, Sirous Ebrahimi, Abbas Alloul, Siegfried E. Vlaeminck, David G. Weissbrodt

## Abstract

Purple non-sulfur bacteria (PNSB), a guild of anoxygenic photomixotrophic organisms, rise interest to capture nutrients from wastewater in mixed-culture bioprocesses. One challenge targets the aggregation of PNSB biomass through gravitational separation from the treated water to facilitate its retention and accumulation, while avoiding the need for membranes. We aimed to produce an enriched, concentrated, well-settling, nutrient-removing PNSB biomass using sequencing batch regimes (SBR) in an anaerobic photobioreactor. The stirred tank was fed with a synthetic influent mimicking loaded municipal wastewater (430-860 mg COD_Ac_ L_Inf_^-1^, COD:N:P ratio of 100:36:4-100:11:2 m/m/m), operated at 30°C and pH 7, and continuously irradiated with infrared (IR) light (>700 nm) at 375 W m^-2^. After inoculation with activated sludge at 0.1 g VSS L^-1^, PNSB were rapidly enriched in a first batch of 24 h: the genus *Rhodobacter* reached 54% of amplicon sequencing read counts. SBR operations at volume exchange ratio of 50% with decreasing hydraulic retention times (48 to 16 h; 1 to 3 cycles d^-1^) and increasing volumetric organic loading rates (0.2 to 1.3 kg COD m^-3^ d^-1^) stimulated the aggregation (compact granules of 50-150 μm), settling (sedimentation G-flux of 4.7 kg h^-1^ m^-2^), and accumulation (as high as 3.8 g VSS L^-1^) of biomass. The sludge retention time (SRT) increased freely from 2.5 to 11 d without controlled sludge wasting. Acetate, ammonium, and orthophosphate were removed simultaneously (up to 96% at a rate of 1.1 kg COD m^-3^ d^-1^, 77% at 113 g N m^-3^ d^-1^, and 73% at 15 g P m^-3^ d^-1^) with a COD:N:P assimilation ratio of 100:6.7:0.9 (m/m/m). Competition for substrate and photons occurred in the PNSB guild. SBR regime shifts sequentially selected for *Rhodobacter* (90%) under shorter SRT and non-limiting acetate concentrations during reaction phases, *Rhodopseudomonas* (70%) under longer SRT and acetate limitation, and *Blastochloris* (10%) under higher biomass concentrations. We highlighted the benefits of a PNSB-based SBR process for biomass accumulation and simultaneous nutrient capture at substantial rates, and its underlying microbial ecology.

**Graphical abstract:** 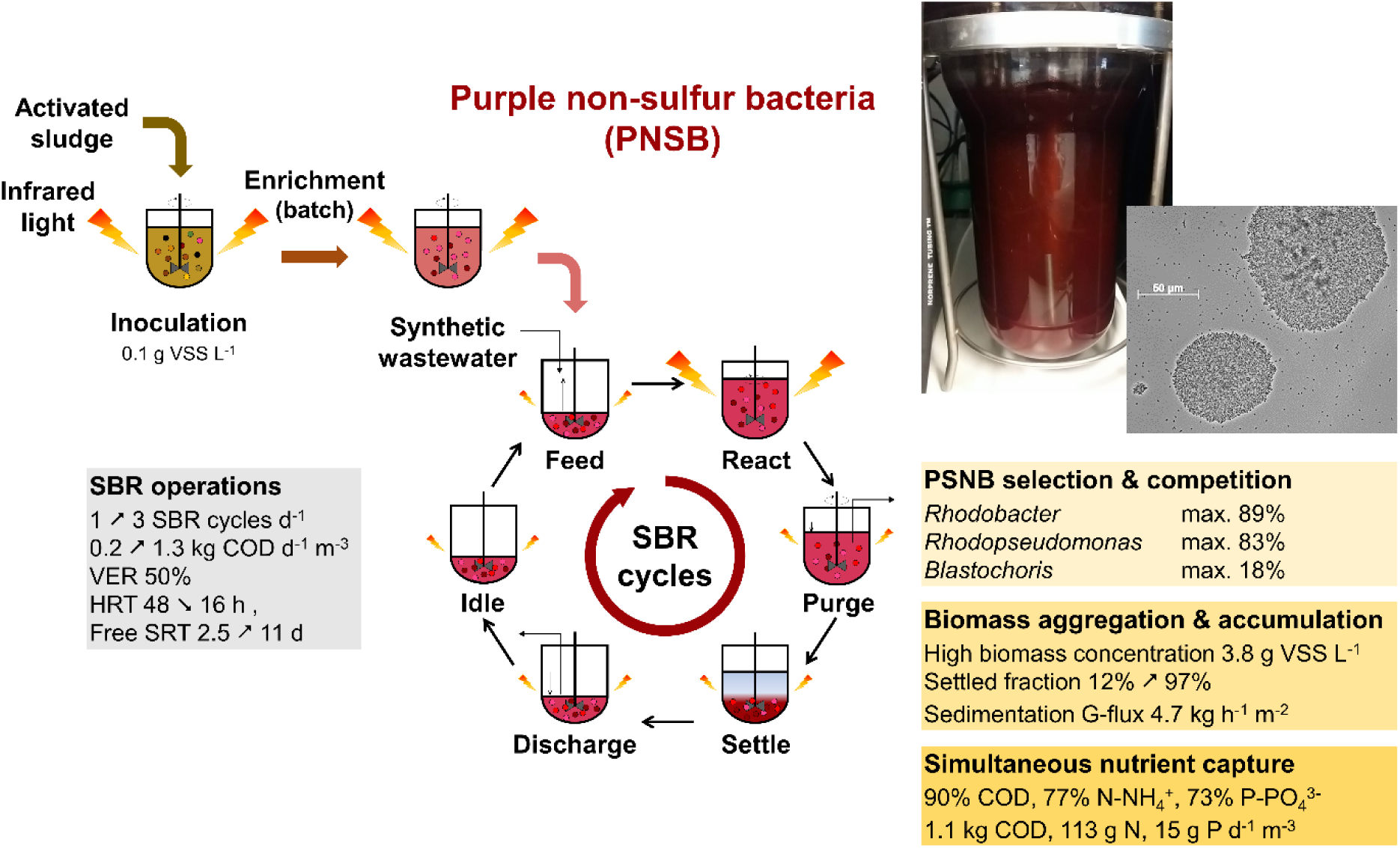

**Highlights:**

- PNSB were highly enriched (90%) in an anaerobic stirred-tank photobioreactor.
- The mixed-culture SBR process fostered PNSB biomass aggregation and accumulation.
- PNSB sludge reached 3.8 g VSS L^-1^ and a sedimentation G-flux of 4.7 kg h^-1^ m^-2^.
- PNSB enabled a high simultaneous removal of COD (96%), N (77%), and P (73%).
- *Rhodobacter*, *Rhodopseudomonas*, and *Blastochloris* competed for acetate and photons.

## 1 Introduction

Biological nutrient removal (BNR) is one of the main goals of wastewater treatment to safeguard aquatic ecosystems from anoxia and eutrophication. Water quality regulations become stricter on the limits of nutrient discharge and removal. European quality criteria target the following residual concentrations and removal of organic matter (125 mg COD_Tot_ L^-1^ and 75% removal; 25 mg BOD_5_ L^-1^ and 70-90% removal), nitrogen (10-15 mg N_Tot_ L^-1^, 70-80% removal), and phosphorus (1-2 mg P_Tot_ L^-1^, 80% removal) in Europe (EUR-Lex, 1991; Guimarães et al., 2018). Besides conventional activated sludge systems, research and innovation target the use of novel microbial processes for water resource recovery (Alloul et al., 2018; Guest et al., 2009; Verstraete and Vlaeminck, 2011) on top of pollution control.

In the resource recovery context, purple non-sulfur bacteria (PNSB) – also referred to purple phototrophic bacteria – can propel a sustainable treatment by capturing nutrient resource from used water (Puyol et al., 2017b; Verstraete et al., 2016), valorizing entropic waste into biomass, bioenergy, bulk chemicals, and biomaterials. PNSB form an attractive guild of anoxygenic, photosynthetic, photoheterotrophic organisms that perform a cyclic photophosphorylation with a facultative anaerobic and hyperversatile metabolism that allows them to grow under ever-changing environmental conditions (Imhoff, 2017; van Niel, 1944). They populate the surface of aquatic environments by accessing and absorbing sunlight energy using carotenoids and bacteriochlorophylls. The spectrum of infrared (IR) electromagnetic wavelengths (800-1200 nm) of light provides them with a competitive advantage in a mixed-culture microbial ecosystem. PNSB can switch between photoorganoheterotrophy, photolithoautotrophy, respiratory or fermentative chemoorgano-heterotrophy, respiratory chemolithoautotrophy, and nitrogen fixation (under ammonium limitations), depending on the composition of electron donors and acceptors present in their surrounding (Madigan and Jung, 2009). This enables them to thrive on different pools of electron donors, recycle electrons, achieve redox homeostasis, and grow under alternation of light and dark (McEwan, 1994). PNSB ferment reduced organics into carboxylates in the dark, photoferment them into dihydrogen, or accumulate and condense them as intracellular storage polymers like biopolyesters (*e.g.*, poly-β-hydroxyalkanoates, PHAs) as electron sinks under nutrient limitations (Hustede et al., 1993). Rediscovering PNSB for ecotechnologies and nutrient capture goes via basic study of their metabolic and selection features from pure to mixed cultures, and eco-design to develop robust, non-axenic, and economically appealing processes (Bryant and Frigaard, 2006).

Their photon-capturing and energy-recycling physiology lead PNSB to achieve rapid biomass specific maximum growth rates (μ_max_) of 1.7 to 5.3 d^-1^ and high biomass yields (Y_X/COD_) on organic substrates (expressed as chemical oxygen demand, COD) of 0.6 to 1.2 g COD_X_ g^-1^ COD_S_ (Eroglu et al., 1999; Hülsen et al., 2014). Electron-balance-wise, this highlights that PNSB can involve additional electron sources from the bulk liquid phase for growth above 1 g COD_X_ g^-1^ COD_S_. Although primarily known as anoxygenic phototrophs, this may happen under conditions that lead PNSB to harness electrons from water molecules like oxygenic photolithoautotrophs. PNSB can assimilate carbon (C), nitrogen (N), and phosphorus (P) from wastewater at COD:N:P ratio of 100:7:2 m/m/m. with an elemental formula for purple phototrophic biomass given as C_1_H_1.8_O_0.38_N_0.18_ (degree of reduction of 4.5 mol e^-^ C-mol^-1^ X_PPB_) (Puyol et al., 2017a). A ratio of 100:5:1 m/m/m and elemental formula of C_1_H_1.8_O_0.5_N_0.2_ has typically been considered for a conventional activated sludge (excl. polyphosphate-accumulating organisms) at a sludge age of 5-8 d (Heijnen and Kleerebezem, 2010; Henze et al., 2000). The potential of PNSB for converting diverse carbon sources such as acetate, malate, butyrate and propionate has been screened with isolates (Alloul *et al*., 2019; Madigan & Gest, 1979a; Stoppani *et al*., 1954), underlying the potential of populations of this guild for water treatment. As phototrophs, PNSB directly use the photonic energy to activate the electrons delivered from electron donors, therefore, maximizing their biomass yield. In contrast, new-generation biological wastewater treatment processes aim to decrease sludge production and handling, by making use of slow-growing and low-yield microorganisms such as polyphosphate-accumulating and anammox bacteria. The use of organisms with a high biomass yield such as PNSB is of definite interest to capture and concentrate carbon, nitrogen and phosphorus nutrient resources out of the wastewater by assimilation into the biomass without need to dissimilate these for energy generation. The produced biomass can be valorized to generate energy through methanization and to produce, *e.g.*, single-cell proteins (*i.e.*, source of microbial proteins), bioplastics via PHAs, and biohydrogen on concentrated streams (Honda et al., 2006; Puyol et al., 2017b).

Technically, one important challenge of photobiotechnologies resides in the limitation of photon supply across the reactor bulk (Pulz O., 2001). Light limitation is often considered *a priori* as a killing factor for the process performance and economics. Therefore, many PNSB-based processes have been operated at concentrations below 1 g VSS L^-1^ to potentially prevent light limitation. Such low biomass concentration can remain a drawback for the intensification of volumetric conversions in the bioprocess.

PNSB have been widely used in pure-culture biotechnologies, *e.g.*, for the production of biohydrogen and biopolymers (Frigaard, 2016; Lenz and Marchessault, 2005; Luongo et al., 2017; Nandi and Sengupta, 1998). Mixed-culture PSNB processes are actively investigated to harness the ability of PNSB to treat wastewater (Hülsen et al., 2016; Nakajima et al., 1997; Verstraete et al., 2016), following the widespread presence and use of these microorganisms in stabilization ponds (Almasi and Pescod, 1996; Freedman et al., 1983). Process configurations involved continuous upflow system (Driessens et al., 1987), continuous-flow stirred tank reactor (Alloul et al., 2019), tubular reactor (Carlozzi et al., 2006), sequencing batch reactor (SBR) (Chitapornpan et al., 2012; Fradinho et al., 2013), membrane bioreactor (MBR) (Hülsen et al., 2016), and membrane sequencing batch reactor (MSBR) (Kaewsuk et al., 2010). One challenge in the application of PNSB organisms is considered to remain in the solid-liquid (S/L) separation of the biomass from the aqueous stream. Decoupling the hydraulic (HRT) and solid (SRT) retention times is crucial to retain the biomass in the process. The use of membrane filtration has been recommended because PNSB have been hypothesized to primarily grow in suspension for catching photons and to settle slowly (Chitapornpan et al., 2012). However, membranes are intended to separate biomass from the treated wastewater, but do not foster the formation of a good settling sludge. In the lab, MBRs are typically used to maintain biomass in suspension (van der Star et al., 2008). A centrifugation step is still needed after the membrane filtration to efficiently concentrate and harvest the PNSB biomass downstream. Thus, alternatives to MBRs can lead to capital and operational savings, since membrane filtration and fouling relate to substantial pumping energy and maintenance costs besides the use of plastic materials. Intensification of PNSB-based environmental biotechnology processes should be targeted by enhancing the bioaggregation and biofilm-forming ability of the biomass. Although previous works have not tailored SBR regimes to this end (Chitapornpan et al., 2012; Fradinho et al., 2013), the application of substrate gradients via SBR operation can be efficient to stimulate microbial aggregation and biomass accumulation, such as typically shown when aiming to granulate activated sludge biomasses (Aqeel et al., 2019; Pronk et al., 2015; Winkler et al., 2018). This should lead to an efficient S/L separation, resulting in lowering costs for downstream processing by potentially reducing the need for ultrafiltration and centrifugation to concentrate the biomass. A SBR design also offers operational flexibility (Morgenroth and Wilderer, 1998) to manipulate reactor cycles and loading rates. Although offering less surface-to-volume ratio, the use of simple stirred-tank designs in SBR application can in addition lead to simpler scale-up than flat-sheet, tubular, or membrane-based processes.

Here, we investigated the possibility to develop a mixed-culture biotechnology process based on the enrichment of a concentrated and well-settling PNSB biomass out of activated sludge in a stirred-tank photobioreactor operated under SBR regime and continuously irradiated with IR light. Conditions to enrich and maintain a PNSB mixed culture were elucidated at bench, along with microbial competition in the PNSB guild. Biomass growth, aggregation, and composition were analysed along with volumetric rates of C-N-P removal. The here-examined microbial ecology insights and aggregation propensity of the PNSB guild can sustain the development of bioengineering strategies for mixed-culture process development in simple SBR design for wastewater treatment and resource recovery from aqueous nutrient streams.

## 2 Material and Methods

### 2.1 Anaerobic, sequencing-batch, stirred-tank photobioreactor setup

The PNSB enrichment was performed in a 1.5-L cylindrical, single-wall, glass, stirred-tank reactor (Applikon Biotechnology, Netherlands) (Figure 1.A). The reactor was inoculated at 0.1 g VSS L^-1^ of flocculent activated sludge taken from the BNR WWTP Harnaschpolder (the Netherlands) after washing the sludge with the cultivation medium 3 times. The reactor operated for 6 months under SBR regime at controlled temperature of 30 ± 1 °C and pH of 7.0 ± 1.0 on an acetate-based synthetic wastewater.

**Figure 1.**
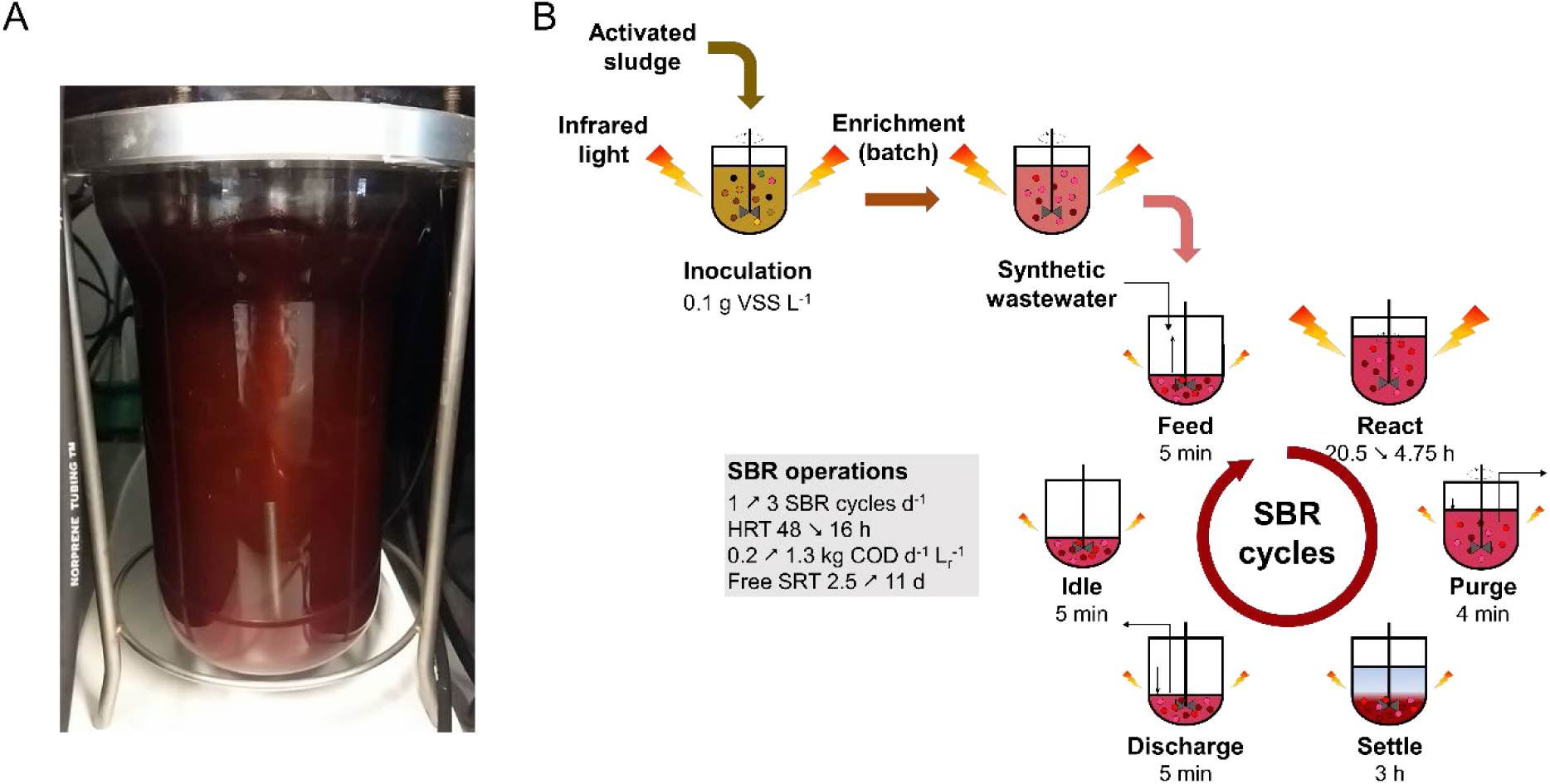
Reactor system and operation **A:** Stirred-tank photobioreactor enriched in PNSB. **B:** PNSB enrichment strategy from activated sludge under SBR regimes. The reactor was inoculated at 0.1 g VSS L^-1^ with BNR activated sludge. One initial batch of 40 h was used to activate the biomass and test the enrichment of PNSB prior to switching to SBR operation over 5 months.

The cultivation medium was calculated based on stoichiometric requirements to sustain PNSB growth and complemented with other minerals adapted from Kaewsuk et al. (2010) to meet with C-N-P anabolic requirements of PNSB. The stock solution consisted of (per liter): 0.914 g of CH_3_COONa·3H_2_O, 0.014 g of KH_2_PO_4_, 0.021 g of K_2_HPO_4_, 0.229 g of NH_4_Cl, 0.200 g of MgSO_4_·7H_2_O, 0.200 g of NaCl, 0.050 g of CaCl_2_·2H_2_O, 0.100 g of yeast extract, 1 mL of vitamin solution, and 1 mL of trace metal solution. The vitamin solution contained (per liter) 200 mg of thiamine–HCl, 500 mg of niacin, 300 mg of ρ-amino-benzoic acid, 100 mg of pyridoxine–HCl, 50 mg of biotin, and 50 mg of vitamin B12. The trace metal solution contained (per liter) 1100 mg of EDTA–2Na·2H_2_O, 2000 mg of FeCl3·6H_2_O, 100 mg of ZnCl2, 64 mg of MnSO_4_·H_2_O, 100 mg of H_3_BO_3_, 100 mg of CoCl_2_·6H_2_O, 24 mg of Na_2_MoO_4_·2H_2_O, 16 mg of CuSO_4_·5H_2_O, 10 mg of NiCl_2_·6H_2_O, and 5 mg of NaSeO_3_. Carbon sources were separated from nitrogen and phosphate sources to avoid contaminations.

To select for purple phototrophs and to avoid the proliferation of green phototrophs, the reactor was placed in a dark fume hood providing only IR light. A white light source was beamed with two halogen lamps (120 W, Breedstraler, GAMMA, Netherlands) placed at the side of the reactor and filtered for IR wavelengths (>700 nm) with two filter sheets (Black Perspex 962, Plasticstockist, UK) placed in front of the lamps. Irradiance was measured at the reactor surface with a pyranometer (CMP3, Kipp & Zonen, Netherlands) and set at a relatively high value of 375 W m^-2^ to promote PNSB enrichment and biomass growth.

After a first 40 h of batch regime to check for the selection of PNSB, the system was switched to an SBR regime, consisting of discharge, idle, feed and settling phases (Figure 1.B). Different cycle timings and HRT, reaction phase length, and COD loading rates were tested as followed in three operational modes. In *SBR1*, 24 cycles of 24 h each were applied (*i.e.*, 24 days of experiment), consisting of: biomass settling (3 h) effluent withdrawal (5 min), influent feeding (5 min), and reaction (20.75 h). In *SBR2*, the total length of the cycles was decreased 3-fold and set at 8 h. The reaction phase was shortened to 4.75 h, while all the other phases were maintained. The reactor was operated over 205 cycles. In *SBR3*, the cycle composition was maintained as in SBR2, while the COD concentration was doubled from 430 to 860 mg COD_Ac_ L^-1^ in the influent to prevent COD-limitations along the reaction phase. All SBRs were operated at a volume exchange ratio of 50%. The stepwise adaptation of the SBR operations from 1 to 3 cycles day^-1^ resulted in HRTs from 48 h (SBR1) to 16 h (SBR2 and SBR3) and in volumetric organic loading rates (OLRs) of 0.215 (SBR1) to 0.645 (SBR2) and 1.290 (SBR3) kg COD d^-1^ m^-3^ (Table 1). The sludge retention time (SRT) was let freely evolve across the SBR operations without controlled purge of the biomass, with median values ranging from 1.5 to 11 d as calculated from eq. 2 in Supplementary material 1, as a result of biomass accumulation.

**Table 1.** Operational conditions of the three SBR regimes across the experimental period. SBR1 was run with a 24-h cycle composed of 20.75 h of reaction time and with 257 ± 54 mg COD L^-1^ in the bulk liquid phase after feeding. In SBR2, the total cycle length was shortened to 8 h, with a reaction time of 4.75 h. In SBR3, the initial COD concentration was doubled compared to the first two SBRs. All SBRs were run at a volume exchange ratio of 50%.

| Process parameters | Units | SBR1 | SBR2 | SBR3 |
| --- | --- | --- | --- | --- |
| **SBR cycles** |  |  |  |  |
| Number of SBR cycles per day | (-) | 1 | 3 | 3 |
| Reaction phase length per cycle | (h) | 20.75 | 4.75 | 4.75 |
| Discharge, idle, feeding phases lengths | (min) | 5 | 5 | 5 |
| Settling phase length | (h) | 3 | 3 | 3 |
| **Retention times** |  |  |  |  |
| Hydraulic retention time (HRT) | (h) | 48 | 16 | 16 |
| Sludge retention time (SRT) <sup>1</sup> | (d) | 1.5 | 7.2 | 10.6 |
| **Loadings** |  |  |  |  |
| Volumetric organic loading rate (OLR) | (g COD d<sup>-1</sup> L<sup>-1</sup>) | 0.215 | 0.645 | 1.29 |
| C:N:P ratio in the influent | (m / m / m) | 100 : 35.8 : 3.8 | 100 : 35.8 : 4.2 | 100 : 11.2 : 1.7 |
| **Measured initial concentrations in the bulk liquid phase at the beginning of reaction phases** |  |  |  |  |
| Acetate | (mg COD L<sup>-1</sup>) | 257 ± 54 | 232 ± 18 | 443 ± 76 |
| Ammonium | (mg N-NH<sub>4</sub><sup>+</sup> L<sup>-1</sup>) | 92 ± 21 | 83 ± 16 | 49 ± 17 |
| Orthophosphate | (mg P-PO<sub>4</sub><sup>3+</sup> L<sup>-1</sup>) | 9.7 ± 1.3 | 9.9 ± 3.3 | 7.7 ± 1.1 |

<sup>1</sup> The sludge retention time (SRT) was let freely evolve over the experimental period. The reactor was operated without purge of biomass. The SRT increased as a result of biomass accumulation in the reactor. The median value over each SBR period is provided. The distributions of SRT are displayed in Figure 2 and detailed evolutions in Supplementary material 3.

The pH of the mixed liquor was controlled at 7.0 ± 1.0 by automatic addition of HCl or NaOH at 1 mol L^-1^ each. The bulk liquid was sparged with argon gas (quality 99.999%) to maintain anaerobic conditions, while continuously stirring at 378 rpm (potentiostat ADI 1012, Applikon, the Netherlands) during the reaction phase.

### 2.2 Analytical methods to measure growth and nutrient consumptions

#### 2.2.1 Measurements of biomass growth and nutrient concentrations

Biomass growth was monitored spectrophotometrically by absorbance at a wavelength of 660 nm (DR3900, Hach, Germany) 4-5 times a week (Supplementary material 2), and gravimetrically by quantifying the concentration of volatile suspended solids (VSS) as described in experimental methods for wastewater treatment (van Loosdrecht et al., 2016). For the 40-batch and SBR1, absorbance measurements were adequate since the biomass was low concentrated and in suspension. For SBR2 and SBR3, the biomass aggregated and VSS measurements were much more accurate.

The consumption of the dissolved nutrients was monitored by sampling the mixed liquor at the beginning and end of the reaction phase, after centrifugation (5 min, 17000 x g) and filtration of the supernatant on 0.45-μm filters (Millex-HV, PVDF, Germany). The concentrations of COD, ammonium (as N-NH_4_^+^) and orthophosphate (as P-PO_4_^3-^) were measured by colorimetric assays (LCK kits no. 114/614/COD, 302/303/ammonium, 348/350/phosphate; Hach-Lange, Dusseldorf, Germany) followed by spectrophotometry (DR3900, Hach, Germany). The COD colorimetric method measured all oxidizable substances (here notably acetate, yeast extract, and EDTA from the trace element solution). As technical control, samples were measured in triplicates and the relative standard deviation was 0.5 - 1.9%.

Acetate concentrations were measured specifically with a high-performance liquid chromatograph (HPLC, Waters, United States) equipped with a Biorad Aminex separation column (HPX-87H, Waters, United States) and an ultraviolet and refraction index (UV/RI) detector (2489, Waters, United States), and using a mobile phase at 1.5 mol H_3_PO_4_ L^-1^ supplied at a flowrate of 0.6 mL min^-1^ and temperature of 60 °C.

#### 2.2.2 Computations of microbial conversions and extraction of growth parameters

All symbols and equations used to compute microbial conversions and extraction of growth parameters are available in Supplementary material 1.

In short, the average percentage of removal (η_S_, %), total rate of nutrient removal (R_S_, kg S d^-1^), apparent volumetric rate of removal of nutrients (r_S_, kg S d^-1^ m^-3^), and apparent growth rate (μ_max_, d^-1^) were calculated using mass balances over the C-N-P nutrients and biomass at a volumetric exchange ratio (VER) of 50%. Measurements of nutrients were performed at the beginning and end of the batch reaction phases of the SBR. Influent concentrations were back-calculated using the VER. The concentrations of nutrients in the effluent were assumed identical as at the end of the reaction phase.

Basic kinetic and stoichiometric parameters for microbial conversions and growth were assessed from nutrient consumptions and biomass production profiles using Aquasim (Reichert, 1994). A mathematical model was constructed using mass balances for substrate consumption and biomass production, and fitted to the experimental data. The maximum biomass-specific rate of acetate consumption (q_S,max_, kg S d^-1^ kg X), maximum yield of biomass production on substrate (Y_X/S,max_, kg X kg^-1^ S), and maintenance rate on substrate (m_S_, kg S d^-1^ kg X) were derived by parameter fit from the Herbert-Pirt relation of substrate allocation for growth and maintenance. The biomass-specific maximum rate of growth (μ_max_, d^-1^) was computed from the relation between q_S,max_ and Y_X/S,max_, assuming the maintenance rate negligible versus the maximum growth rate during the exponential phase of the batch reaction period.

### 2.3 Analysis of biomass and microbial community compositions

#### 2.3.1 Light microscopy analysis of microbial morphotypes and bioaggregates

Microbial morphotypes present in the enrichment were visually observed by phase contrast microscopy (Axioplan 2, Zeiss, Germany).

#### 2.3.2 Wavelength scan analysis of pigment content in the PNSB-enriched biomass

The evolution of the biomass contents in bacteriochlorophyll a (BChl a) and carotenoids in the biomass was used as a proxy for tracking the PNSB enrichment in the mixed liquor. Measurements were performed by wavelength scan over the visible and near-infrared spectrum from 400 to 1000 nm (DR3900, Hach, Germany). A focus was attributed to absorbance peaks between 800-900 nm (BChl a) and 400-600 nm (carotenoids).

#### 2.3.3 V3-V4 16S rRNA gene amplicon sequencing of bacterial community compositions

Genomic DNA was extracted from biomass samples throughout the duration of the experiment, using UltraClean Microbial Isolation kits (MOBIO laboratories, Inc., USA) following manufacturer’s instructions, and stored at -20 °C. The concentrations and qualities of the DNA extracts were measured by Qbit3 fluorimeter (Thermofisher Scientific, USA), according to manufacturer’s instructions. The DNA extracts were sent to Novogene (China) for amplicon sequencing. The V3-V4 regions of the 16S rRNA gene were amplified by polymerase chain reaction (PCR) using the set of forward V3-V4 forward 341f (5’-CCTACGGGAGGCAGCAG-3’) and reverse 806r (5’-GGACTACHVGGGTWTCTAAT-3’) primers (Takahashi *et al*., 2014). The amplicon sequencing libraries were pooled and sequenced in an Illumina paired-end platform. After sequencing, the raw read were quality filtered, chimeric sequences were removed and OTUs were generated on the base of ≥ 97% identity. Subsequently, microbial community analysis was performed by Novogene using Mothur & Qiime software (V1.7.0). For phylogenetical determination the most recent SSURef database from SILVA (http://www.arb-silva.de/) was used. Relative abundances of OTUs were reported as % total sequencing reads count.

## 3 Results

### 3.1 High, simultaneous nutrient removal was achieved in the PNSB-enriched SBR

The nutrient removal performances achieved by the PNSB-based process from SBR1 to SBR2 and SBR3 regimes are displayed in figure 2 and Table 2. The detailed dynamics in nutrient and biomass concentrations and compositions are provided in Supplementary material 3.

**Figure 2.**
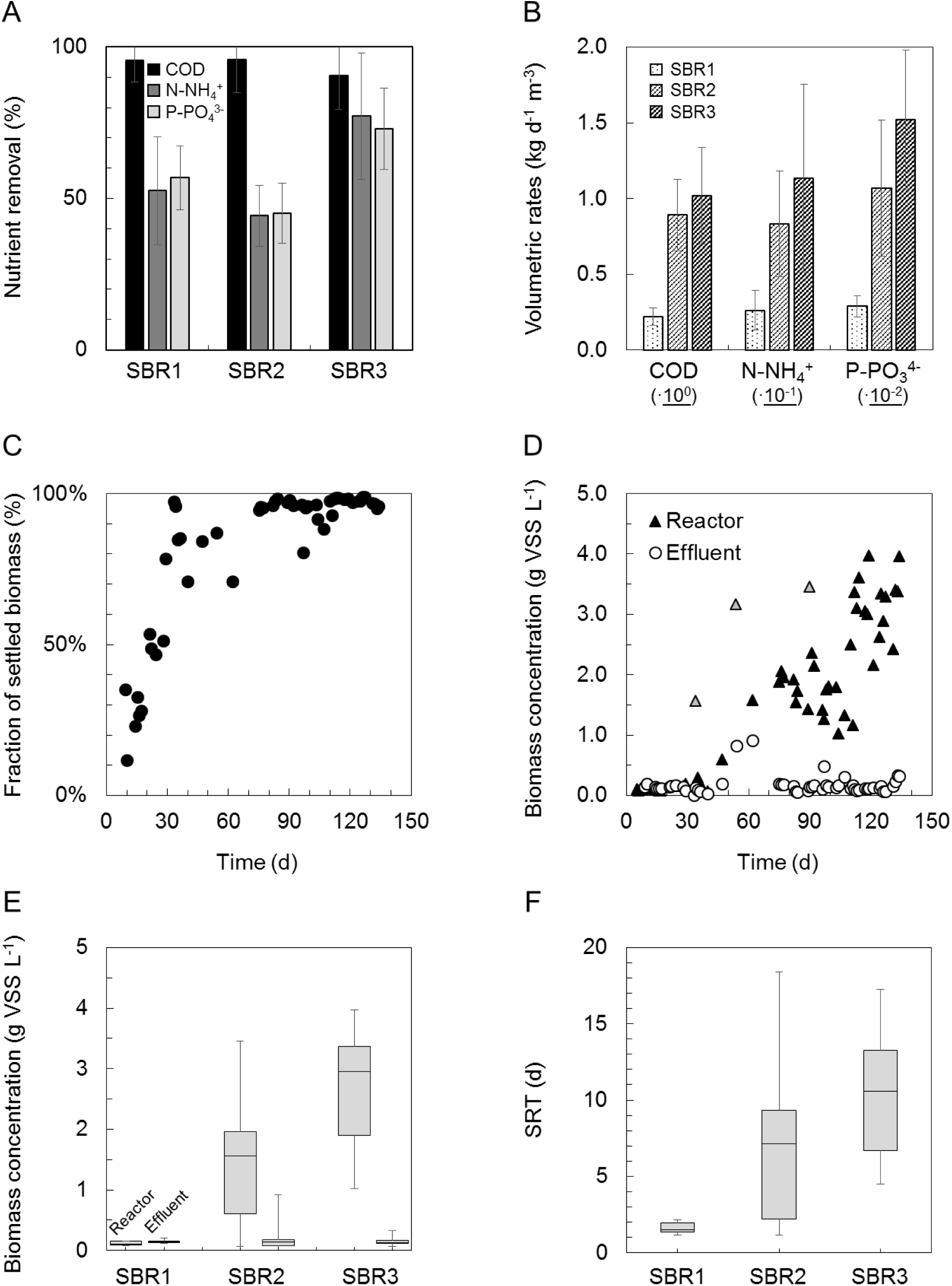
Nutrient removal and biomass characteristics across SBR operations in the mixed-culture PNSB photobioreactor. **A:** Increases in COD, ammonium, and orthophosphate nutrient removal percentages from SBR1 to SBR3. On average, 95% of the COD was removed during all operational states. N-NH_4_^+^ and P-PO_4_^3-^ reached 77% and 73% of removal from the synthetic influent. **B:** Gradual increases in volumetric rates of C-N-P nutrient removals from SBR1 to SBR3. **C:** Increase in the fraction of mixed-liquor biomass that settled in the bioreactor along SBR1 (days 3-30), SBR2 (days 30-100), and SBR3 (days 100-135) after inoculation with BNR activated sludge and a first batch of 40 h. **D:** Accumulation of biomass in the photobioreactor from SBR1 to SBR2 and SBR3. The grey triangles relate to cleanings and resuspensions of the wall biofilm in the biosystem. They indicate the total amount of biomass that accumulated in the reactor. **E:** Distributions of biomass concentrations in the reactor at the end of the reaction phase and in the effluent after settling. The settling ability of the biomass increased steadily during SBR operations. The residual biomass concentration in suspension at the end of the settling phase in SBR3 was 10 times lower than the concentration in the mixed liquor during reaction time, displaying the well-settling property of the PNSB-enriched biomass. **F:** The SRT was let freely evolve in the reactor, increasing from median values of 2 d (SBR1) to 7.5 d (SBR2) and 11 d (SBR3) along with biomass accumulation.

**Table 2.**
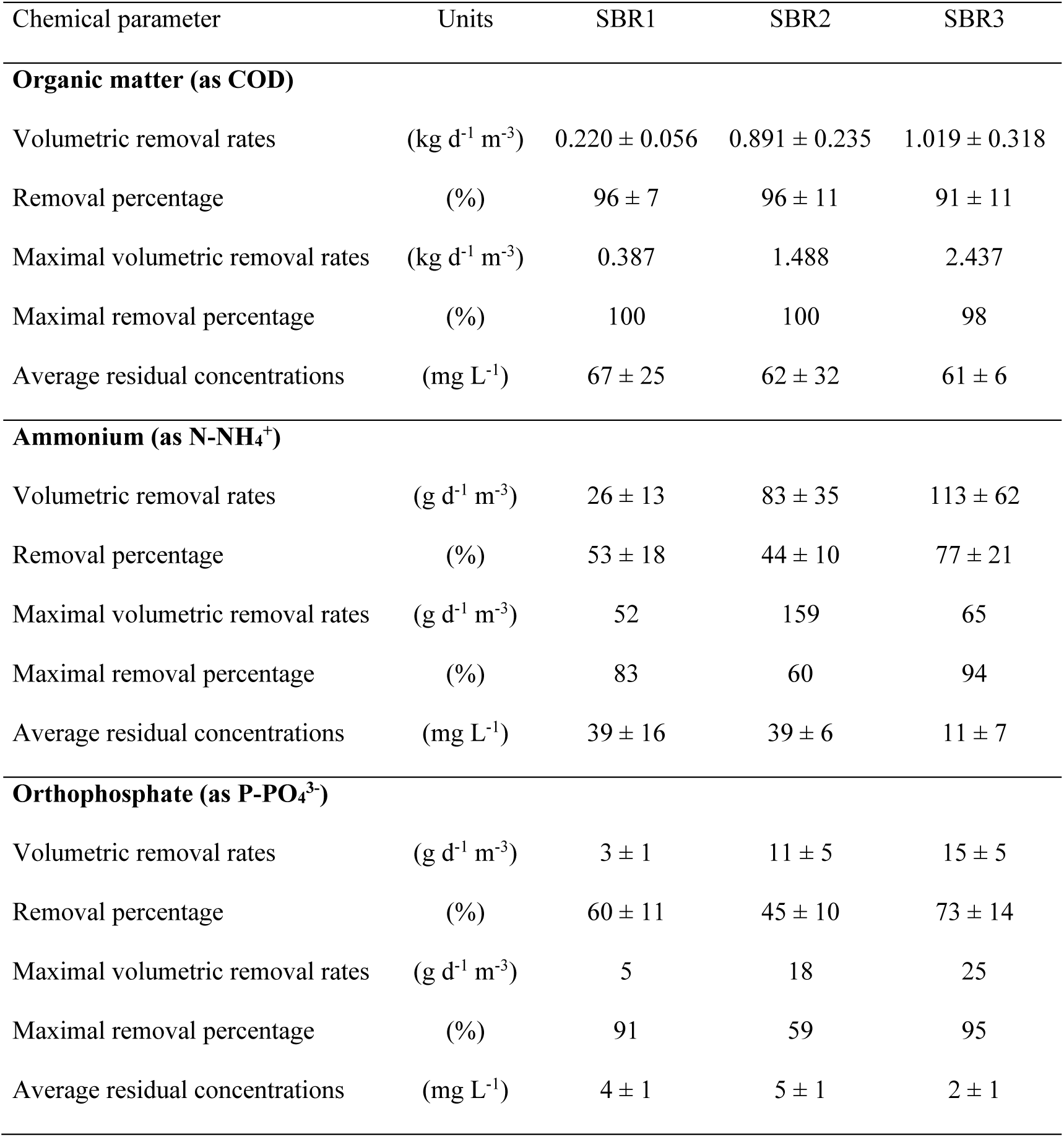
Nutrient removal by the PNSB-enriched biomass in SBR1, SBR2, and SBR3 presented as averages and maximal values of removal rates and removal percentages. Residual concentration and removal percentages for COD met with European legislation limits for all SBRs (averages above 90% removal, with residues close to 60 mg COD L^-1^). Ammonium and orthophosphate were removed up to 77% and 73%, respectively, with residual concentrations reaching 11 mg N-NH_4_^+^ L^-1^ and 2 mg P-PO_4_^3-^ L^-1^.

| Chemical parameter | Units | SBR1 | SBR2 | SBR3 |
| --- | --- | --- | --- | --- |
| **Organic matter (as COD)** |  |  |  |  |
| Volumetric removal rates | (kg d<sup>-1</sup> m<sup>-3</sup>) | 0.220 ± 0.056 | 0.891 ± 0.235 | 1.019 ± 0.318 |
| Removal percentage | (%) | 96 ± 7 | 96 ± 11 | 91 ± 11 |
| Maximal volumetric removal rates | (kg d<sup>-1</sup> m<sup>-3</sup>) | 0.387 | 1.488 | 2.437 |
| Maximal removal percentage | (%) | 100 | 100 | 98 |
| Average residual concentrations | (mg L<sup>-1</sup>) | 67 ± 25 | 62 ± 32 | 61 ± 6 |
| **Ammonium (as N-NH<sub>4</sub><sup>+</sup>)** |  |  |  |  |
| Volumetric removal rates | (g d<sup>-1</sup> m<sup>-3</sup>) | 26 ± 13 | 83 ± 35 | 113 ± 62 |
| Removal percentage | (%) | 53 ± 18 | 44 ± 10 | 77 ± 21 |
| Maximal volumetric removal rates | (g d<sup>-1</sup> m<sup>-3</sup>) | 52 | 159 | 65 |
| Maximal removal percentage | (%) | 83 | 60 | 94 |
| Average residual concentrations | (mg L<sup>-1</sup>) | 39 ± 16 | 39 ± 6 | 11 ± 7 |
| **Orthophosphate (as P-PO<sub>4</sub><sup>3-</sup>)** |  |  |  |  |
| Volumetric removal rates | (g d<sup>-1</sup> m<sup>-3</sup>) | 3 ± 1 | 11 ± 5 | 15 ± 5 |
| Removal percentage | (%) | 60 ± 11 | 45 ± 10 | 73 ± 14 |
| Maximal volumetric removal rates | (g d<sup>-1</sup> m<sup>-3</sup>) | 5 | 18 | 25 |
| Maximal removal percentage | (%) | 91 | 59 | 95 |
| Average residual concentrations | (mg L<sup>-1</sup>) | 4 ± 1 | 5 ± 1 | 2 ± 1 |

#### 3.1.1 Nutrient removing activities were detected during the initial batch

During the first 40 h of batch used to activate the biomass, nutrients were removed at 98 % COD, 52 % N-NH_4_^+^, and 60% P-PO_4_^3-^ (Figure 2.A). These related to apparent volumetric removal rates of 0.190 ± 0.048 kg COD d^-1^ m^-3^, 21.5 ± 8.6 g N d^-1^ m^-3^, and 2.5 ± 0.5 mg P d^-1^ m^3^ (Figure 2.B). The COD:N:P consumption ratio was 100:7.5:0.12 (m/m/m) in this batch.

#### 3.1.2 A complete removal of acetate was achieved across all SBR operation modes

During SBR1, the average percentage of removal of the acetate-based biodegradable COD was 96%, with an average volumetric consumption rate of 0.220 ± 0.060 kg COD d^-1^ m^-3^. During SBR2, 96% of the COD was removed as well at a 4-fold higher rate of 0.891 ± 0.235 kg COD d^-1^ m^-3^. The carbon was fully removed over the first hour of the reaction phase, resulting in remaining 3.75 h of substrate limitation. During SBR3, the COD load in the influent was doubled, and as a result, no nutrient limitation occurred during the reaction phase. COD remained highly removed at 91%, with a volumetric removal rate of 1.08 ± 0.32 kg d^-1^ m^-3^.

#### 3.1.3 A maximum of 85% of ammonium and 74% of phosphate was removed from the inflow

The ammonium removal rates increased from 26 ± 13 g N-NH_4_^+^ d^-1^ m^-3^ in SBR1 to 83.4 ± 35 g N d^-1^ m^-3^ during SBR2, and 113.3 ± 62 g N d^-1^ m^-3^ during SBR3. Average N-removal percentages evolved from 53% to 44% and 77% of the ammonium load across the three SBRs, respectively. Removal rates of orthophosphate increased from 3.0 ± 0.7 g P-PO_4_^3-^ d^-1^ m^-3^ of SBR1 to 10.7 ± 4.5 g P d^-1^ m^-3^ in SBR2 and 15.2 ± 4.6 g P d^-1^ m^-3^ in SBR3, with average P-removal percentages of 57, 45 and 73% per cycle, respectively. Under the non-limiting COD conditions of SBR3, the acetate, ammonium, and orthophosphate were released at median concentrations of 43 (min = 17; 1^st^-3^rd^ quartile = 28-62) mg COD L_Eff_^-1^, 10 (2; 8-13) mg N-NH_4_^+^ L_Eff_^-1^, and 2.0 (0.4; 1.7-2.9) mg P-PO_4_^3-^ L_Eff_^-1^, *i.e.*, close to European discharge criteria.

Thus, the average apparent COD:N:P assimilation ratio evolved from 100:7.5:0.12 in the batch to 100:9.2:1.2 (SBR1-2) under COD-limitation and 100:6.7:0.9 (SBR3) under non-COD-limitation. Across and beyond the experimental period, the intrinsic kinetics and stoichiometry of the PNSB-enriched biomass ranged with a biomass specific maximum growth rate (μ_max_) of 0.96-2.16 d^-1^ and a maximum yield of biomass production on substrate (Y_X/COD,max_) of 0.21-0.74 g VSS g^-1^ CODs, respectively. This related to a yield value of 0.34-1.19 g CODx g^-1^ CODs when using a theoretical elemental composition of C_1_H_1.8_O_0.38_N_0.18_ (1.607 g CODx g^-1^ VSS) for purple phototrophic bacteria (Puyol et al., 2017a). The maximum biomass specific consumption of acetate (q_COD,max_) ranged from 0.03-0.78 kg CODs h^-1^ kg^-1^ VSS. These measurements were performed directly during SBR cycles at the actual concentration of the biomass present in the system. More accurate measurements and derivation of these physiological parameters can be performed at diluted initial concentrations of PNSB biomass to prevent nutrient and light limitations during batch tests.

#### 3.1.4 Kinetic and stoichiometric parameters of microbial growth of the PNSB mixed culture

The maximum biomass specific rate of substrate consumption (q_S,max_), the yield of biomass growth of substrate consumption (Y_X/S_) and the maximum biomass specific growth rate (μ_max_) were obtained by parameter fit to batch evolutions of acetate and biomass during selected reaction phases of the SBRs (Table 3). Under the conditions of the initial batch and of SBR1 (*i.e.*, long HRT of 48 h, low OLR of 0.215 kg COD d^-1^ m^-3^, very low biomass concentration of 0.1 g VSS L^-1^, and low SRT of 1.5 d), the highly-enriched PNSB biomass displayed a high q_S,max_ between 5.8-13.7 g COD_S_ d^-1^ g^-1^ VSS on which it maximized its growth rate: μ_max_ (2.16-3.36 d^-1^) was substantial, ranging between values reported in literature for mixed cultures and pure cultures of PNSB. The biomass thus developed at a relatively low yield Y_X/S, max_ (0.23-0.39 g VSS g^-1^ COD_S_). The maintenance rate (m_s_) was estimated to 0.72 g COD_S_ d^-1^ g^-1^ VSS. Under the conditions of SBR2 and SBR3 (*i.e.*, 3-times lower HRTs, 3 to 6-times higher OLRs, 16 to 30-times higher biomass concentrations, and 5 to 7-times longer SRTs), the biomass consumed acetate at a 3 to 8-fold lower q_S,max_ (1.8-2.2 g COD_S_ d^-1^ g^-1^ VSS). The maximum growth rate and yield values could not be extracted from the data collected from the reactions phases at high biomass concentration (low sensitivity of absorbance and VSS measurements to detect changes). More accurate estimates are obtained with batch tests conducted with a diluted concentration of biomass.

**Table 3.** Observed physiological parameters of the biomass of the PNSB mixed culture extracted from the reaction periods of the initial batch and the three SBR periods, and comparison with literature data obtained from pure-culture and mixed-culture PNSB systems. The values are given based on measured COD units for the acetate substrate and absorbance-calibrated VSS units for the biomass. The elemental formula for purple phototrophic bacteria C_1_H_1.8_O_0.38_N_0.18_ (4.5 mol e^-^ C-mol^-1^, 36 g COD C-mol^-1^, 22.4 g VSS C-mol^-1^, 1.607 g COD_X_ g^-1^ VSS) (Puyol et al., 2017a) may be used for conversion of VSS in to COD units.

| System | $q_{S,\max}$<br>($\text{g COD}_S \text{ d}^{-1} \text{ g}^{-1} \text{ VSS}$) | $Y_{X/S,\max}$<br>($\text{g VSS g}^{-1} \text{ COD}_S$) | $\mu_{\max}$<br>($\text{d}^{-1}$) |
| --- | --- | --- | --- |
| PNSB pure cultures | n.a. | 0.98-1.23 <sup>a</sup> | 5.28 <sup>a</sup> |
| PNSB mixed culture | n.a. | 0.23-0.63 <sup>b</sup> | 0.72-1.68 <sup>b</sup> |
| Initial batch | 5.76 | 0.39 | 2.16 |
| SBR1 <sup>c</sup> | $13.68 \pm 5.04$ | $0.23 \pm 0.02$ | $3.36 \pm 1.4$ |
| SBR2 <sup>c</sup> | $1.76 \pm 0.91$ | n.a. <sup>d</sup> | n.a. <sup>d</sup> |
| SBR3 <sup>c</sup> | $2.22 \pm 0.79$ | n.a. <sup>d</sup> | n.a. <sup>d</sup> |

<sup>a</sup> Taken from literature (Eroglu et al., 1999; Jih, 1998)
<sup>b</sup> Taken from literature (Hülßen et al., 2016; Hülßen et al., 2014; Kaewsuk et al., 2010; Puyol et al., 2017)
<sup>c</sup> Average values collected over different cycles monitored over SBR1 (cycles 11, 17), SBR2 (cycles 21, 145, 166), and SBR3 (cycles 10, 31, 49, 91).
<sup>d</sup> High biomass concentrations in the system. VSS and absorbance measurement are no sensitive enough to detect growth over reaction period. External batches with diluted biomass should be performed to this end.

### 3.2 SBR operations enhanced the settling ability and accumulation of the PNSB biomass

The settling ability of the PNSB biomass increased across enrichment SBR operations, leading to substantial accumulation of biomass in the system from 0.1 (SBR1) to 1.6 (SBR2) to 3.0 (SBR3) g VSS L^-1^ as median values (Figure 2.C-D). The enhancement of the settling ability was measured by comparing these biomass concentrations present in the mixed liquor at the end of the reaction phase with the concentrations in the effluent after the settling phase which was for all SBRs as low as 0.13-0.15 g VSS L_Eff_^-1^ (Figure 2.C-D). The fraction of settled biomass increased across SBR1 from 12% to 53% of the VSS present in the mixed liquors at the end of reaction phases, reached 96% by end of SBR2, and remained high at 97±3% over SBR3 (Figure 2.C). The total rates of biomass accumulation calculated over the full settling period of 3 h increased from 0.02±0.01 (SBR1) to 0.69±0.46 (SBR2) and 1.30±0.45 (SBR3) g VSS h^-1^, or from 0.02±0.01 to 0.46±0.31 and 0.87±0.30 kg VSS h^-1^ m^-3^, respectively, when translated into volumetric rates. At the beginning of SBR1, the full 3 h period was required to settle the suspended biomass. At the end of SBR3, most of the 5.9 g VSS of biomass that aggregated and accumulated in the system settled in about 10 min (*i.e.*, 35 gVSS h^-1^ or 24 kg VSS h^-1^ m^-3^ effectively). This high settling rate obtained on SBR3 corresponds to a sedimentation G-flux of solids of 4.7 kg h^-1^ m^-2^. This displayed the well-settling property of the aggregated PNSB biomass. It underlined potential for considerably shortening the settling phase and SBR cycle length in order to increase the daily loading of the system.

The fraction of VSS in the TSS remained relatively high with 85% (SBR1) to 93% (SBR2) to 80% (SBR3) as median values, *i.e.*, corresponding to a fraction of inorganic suspended solids (ISS) between 7-20%. During SBR3 a period at lower VSS fraction with values below 60% and higher ISS fraction (>40%) was detected between days 97-117, underlying potential accumulation of inorganics, *e.g.*, as intracellular polyphosphate (not measured), during nutrient assimilation in the biomass. Taking into account the P-uptake (average = 9, max = 13 mg P-PO_4_^3-^ cycle^-1^), biomass production of 90 mg VSS cycle^-1^ (using the measured yield of 0.23±0.02 g VSS g^-1^ CODs) and VSS fraction of 80-90%, the average and maximum P-content of the cell were calculated to 8 and 14% of the cell dry weight which can underlie intracellular storage of polyphosphate to some extent.

The SRT was let to increase freely, without controlled purge of biomass, as a result of the enhancement of settling properties of the biomass: it rose from 2 d in SBR1 to 7 d in SBR2 and 11 d in SBR3 as median values (Figure 2.D). Strategies can be tested to control the SRT at specific values on the range between, *e.g.*, 3-10 days, depending on nutrient capture and biomass production targets.

### 3.3 Microscopy images displayed an increasing size of compact granular bioaggregates

The PNSB enrichment process could be easily be tracked visually with the gradual increase in the purple color intensity in the bioreactor (Figure 3.A-D).

**Figure 3.**
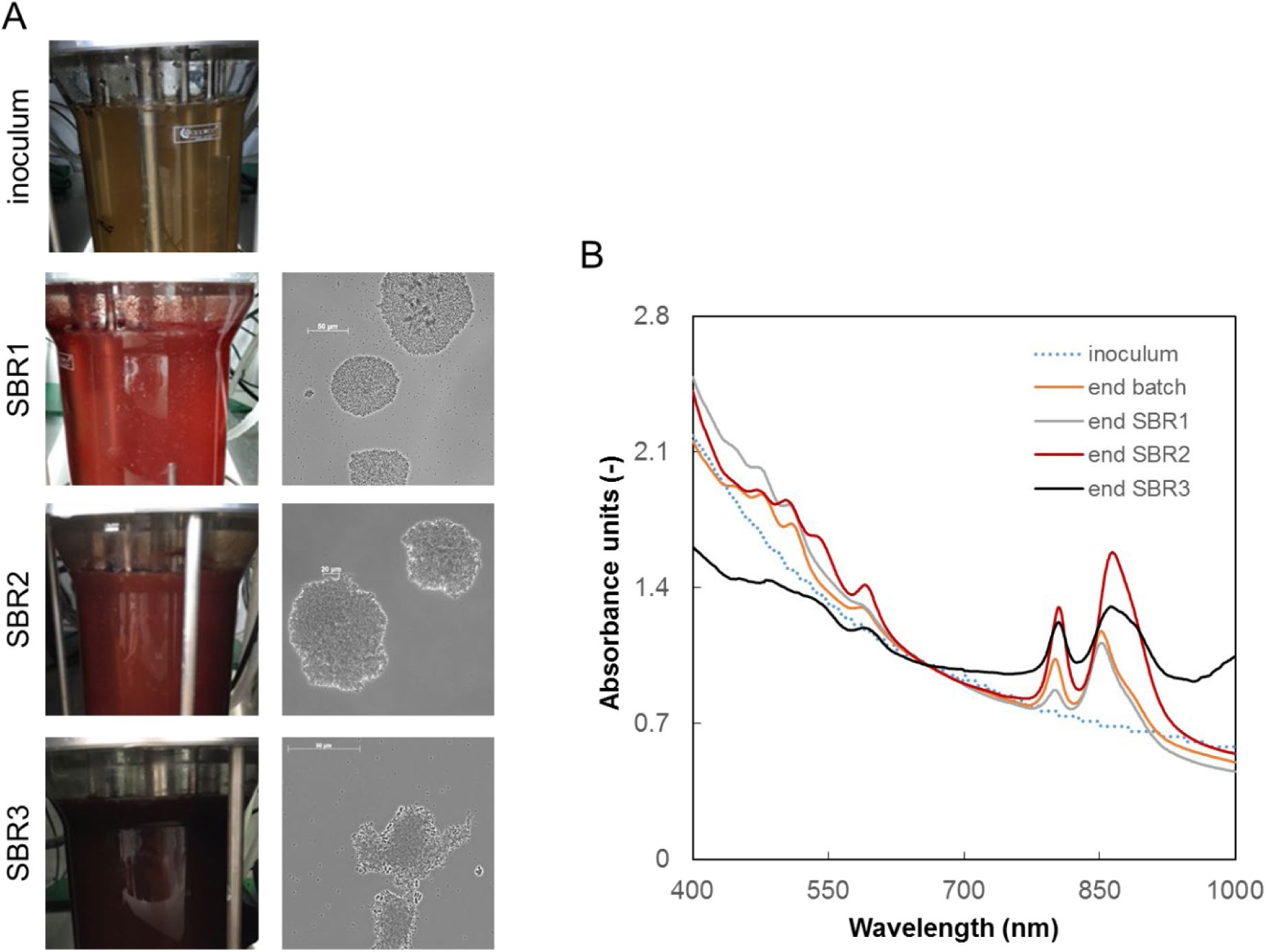
Evolution of the pigmentation and aggregative characteristics of the PNSB-enriched biomass. **A:** Pictures of the PNSB-enriched biomass taken with a digital camera and phase-contrast microscopy images of the aggregates present in SBR1 to SBR3. The biomass in the reactor changed from brown to purple-red colour during the enrichment after inoculation with activated sludge. The size of the aggregates increased during time along with increased settling abilities of the biomass. **B:** Wavelength scans of intact cultures, normalized for the biomass content (at 660 nm). The presence of PNSB was tracked at peaks around 800-900 nm (coding for Bchl a) and 400-500 nm (carotenoids). After the initial batch phase of 40 h, the peaks typical for PNSB pigments were present, and persisted in the biomass until the end of SBR3.

After inoculation with flocculent activated sludge, phase-contrast microscopy imaging revealed the presence of dense aggregates already in SBR1 formed by the PNSB biomass (Figure 3.E-G). Some cells clustered in flower-shaped aggregates, in a way comparable to the typical morphotype of *Rhodopseudomonas*. Other rod-shaped cells were present, putatively belonging to *Rhodobacter* and *Blastochloris* genera. The size of the aggregates increased from 50 to 150 μm during the operational time along with the better settling abilities of the biomass.

### 3.4 Wavelength scans highlighted the enrichment of carotenoid and bacteriochlorophyll pigments in the biomass and a shift in PNSB populations

Carotenoids and bacteriochlorophylls, and their increase along the enrichment of the PNSB biomass, were detected by the presence of absorbance peaks at wavelengths between 450-500 nm and between 800-900 nm. The wavelength scan data presented in Figure 3.H are normalized by the biomass concentrations, expressed as absorbance units at 660 nm. Peaks at 800 nm and 850 nm were already present at the end of the initial batch phase, and persisted during SBR1. At the end of SBR2, the absorbance peaks shifted to higher wavelengths of 805 nm and 865 nm. During SBR3, another peak was detected at 1000 nm that is characteristic for the genus *Blastochloris*. This suggested a shift in predominant microbial populations harbouring different types of pigments in the PNSB guild across the mixed-culture enrichment process.

### 3.5 Amplicon sequencing revealed selection shifts from *Rhodobacter* to *Rhodopseudomonas* and *Blastochloris* genera within the guild of PNSB

The composition of the bacterial community of the mixed culture and underlying shifts in predominant populations were analysed by V3-V4 16S rRNA gene amplicon sequencing. The times series of PNSB populations are displayed in Figure 4. The detailed times series of the full set of identified genera in the sequencing dataset is given in Supplementary material 5.

**Figure 4.**
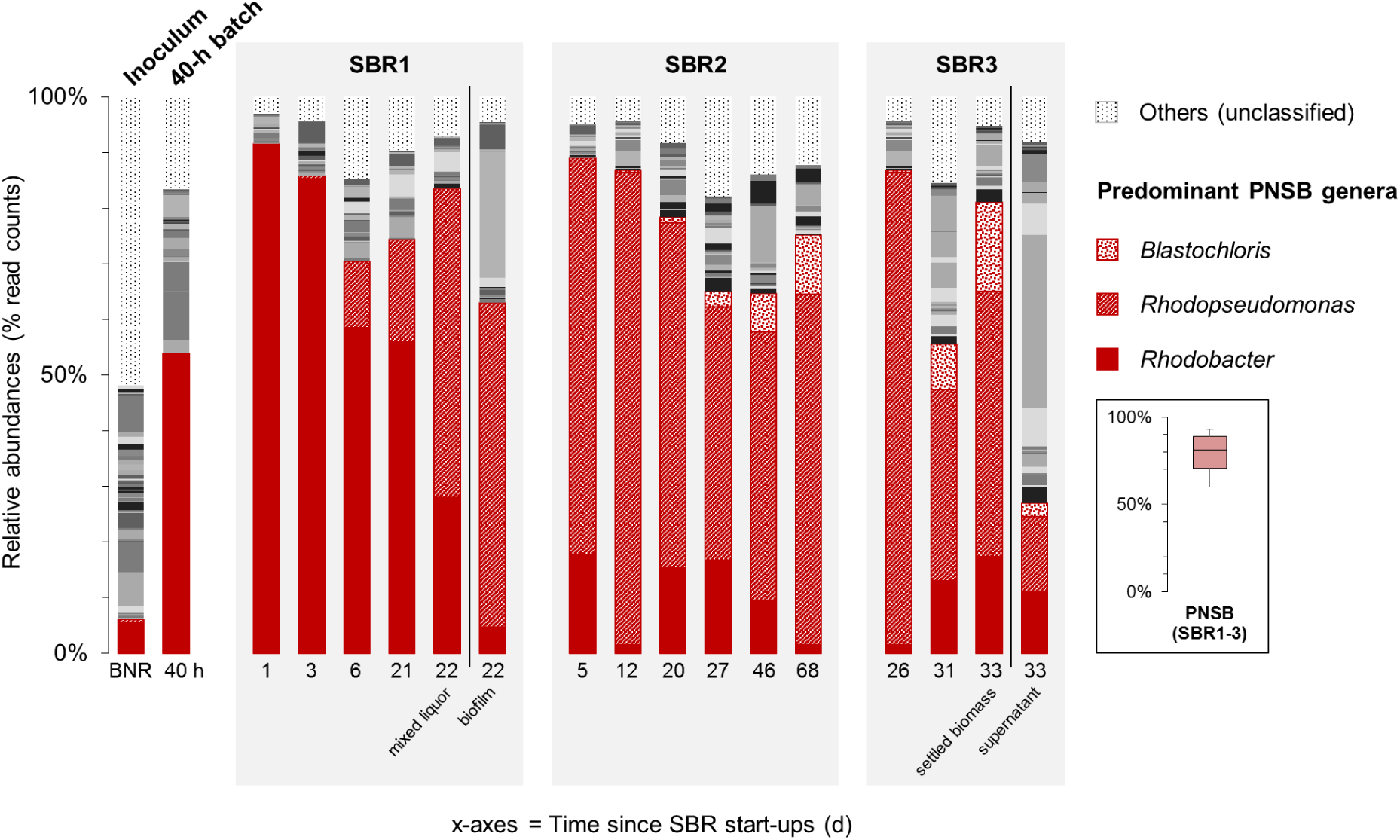
Time series of V3-V4 16S rRNA gene amplicon sequencing of bacterial community compositions in the PNSB-enriched mixed-culture process along SBR regime shifts. After inoculating the reactor with BNR activated sludge (“BNR”), a first PNSB genus *Rhodobacter* was initially enriched during the first 40-h batch (“40 h”) and early SBR1 period. The second PNSB genus *Rhodopseudomonas* was predominantly selected across operations of SBR2 and SBR3. The third PNSB genus *Blastochloris* popped up by end of SBR2 and SBR3. The PNSB guild remained predominant in the biomass across the process with an average total relative abundance of sequencing reads affiliated to known PNSB above 60% of the total community dataset (median = 81%; min-max = 60-93%). In SBR1, both the mixed liquor and the wall biofilm were sampled on day 22 and sequenced. In SBR3, both the settled biomass and s were sampled on day 33 after settling, and sequenced. The full set of genera is given in Supplementary material 5.

Across the experimental period, a rapid initial selection for a low number of predominant organisms occurred under the conditions of the first batch (acetate-based synthetic feed, non-limiting IR irradiance), prior to establishment of a more diverse community along SBR cycles. The BNR activated sludge inoculum presented a diversity of genera, with *Rhodobacter* as the main PNSB detected at 5% of the sequencing read counts. The typical populations of the BNR sludge like ammonium oxidizer (*Nitrosomonas*), nitrite oxidizer (*Nitrospira*), denitrifier (*Zoogloea*), polyphosphate-(“*Candidatus* Accumulibacter”) and glycogen-accumulating (“*Ca.* Competibacter”) organisms got rapidely outcompeted right after start-up of the first batch under PNSB-selective conditions (Supplementary material 5).

At the end of the 40-h batch phase, *Rhodobacter* reached a relative abundance of 52%. At the end of the first cycle of SBR1, a high-grade enrichment of 90% of *Rhodobacter* was obtained. Around the 10^th^ cycle of SBR1 (10 days), the genus *Rhodopseudomonas* got enriched at 15%, and reached 50% at the end of the 23^rd^ cycle (23 days after inoculum). The compositions of the communities of the mixed liquor and of the biofilm that developed on the walls of the reactor during the 13^th^ cycle revealed that *Rhodopseudomonas* (55%) was outcompeting *Rhodobacter* (5%) in the biofilm, while *Rhodobacter* (60%) was more enriched than *Rhodopseudomonas* (10%) in the mixed liquor. Then, *Rhodobacter* decreased constantly from cycle to cycle, while *Rhodopseudomonas* progressively took the lead in the flocculent biomass as well.

After 18 cycles of SBR2 (*i.e.*, 4.5 and 22.5 days from the starts of SBR2 and SBR1, respectively), *Rhodopseudomonas* became dominant (70%), outcompeting *Rhodobacter* (17%) in the enrichment culture. Interestingly, after 20 days of SBR2, the genus *Blastochloris*, also an affiliate of the PNSB guild, got selected, while the relative abundance of *Rhodopseudomonas* decreased to 60% at the end of SBR2. In SBR3, *Blastochloris* reached 10% of the bacterial community dataset.

## 4 Discussion

### 4.1 A high-grade enrichment of a concentrated, well-settling PNSB biomass was obtained under SBR regime

The enrichment of PNSB has often been successful, while most PNSB mixed cultures reported so far have mainly been in membrane systems (Hülsen et al., 2016). Here, we successfully enriched a mixed culture of PNSB out of activated sludge under traditional SBR regime in a stirred-tank system without the use of a membrane module to separate the biomass and the bulk liquid phase. This went by using the natural propensity of PNSB to form biofilms and bioaggregates. SBR regimes result in substrate gradients across reactor operation from high concentrations at the beginning of the cycle to low residual concentrations at the end. Such substrate gradients are known to promote the bioaggregation of microorganisms (Aqeel et al., 2019; Pronk et al., 2015; Winkler et al., 2018).

Promotion of bioaggregation of PNSB is key for a good S/L separation and accumulation of biomass in the system. One important outcome of this study highlighted that aggregation of PNSB can be stimulated under SBR regime to intensify the volumetric conversions and to facilitate downstream processing. After inoculation at 0.1 g VSS L^-1^, a high concentration of a PNSB-enriched biomass of up to a maximum of 4.0 g VSS L^-1^ was obtained in SBR3. The good settling ability of the PNSB biomass obtained under this regime resulted in the emission of less than 5% of the mixed liquor biomass in the effluent of SBR3, as low as 0.1 g VSS L^-1^. Interestingly, Driessens et al. (1987) have early reported the flocculation and good sedimentation (G-flux of 7-9 kg h^-1^ m^-2^ comparable to well-flocculated activated sludge) of *Rhodobacter capsulatus* in an upflow continuous photobioreactor operated under loading rates of 2.5-5.0 kg C d^-1^ m^-3^ (as calcium lactate; *i.e.*, 6.7-13.3 kg COD d^-1^ m^-3^) and 0.5-1.0 kg N d^-1^ m^-3^ (as ammonium) with 87% C and N assimilation in the biomass (3.3-4.2 g VSS L^-1^). The PNSB-enriched biomass during SBR3 displayed a high sedimentation G-flux of 4.7 kg h^-1^ m^-2^ relatively close to the values reached by Driessens et al. (1987) under highly concentrated loading rates 5 to 10-fold higher than used here (max. 1.3 kg COD d^-1^ m^-3^ in SBR3). Collectively, this comparison sustains that PNSB can be aggregated for a higher accumulation and retention of biomass to intensify nutrient conversions.

In the PNSB mixed culture, the HRT was initially set high to 48 h (*i.e.*, 1 cycle d^-1^ at a volume exchange ratio of 50%) to maintain biomass during start-up, prior to decreasing it to 16 h (3 cycles d^-1^) from SBR2 onward. This value was in the range of the HRTs of 8-24 h that have been used in the operation of continuous photo anaerobic membrane bioreactor (PAnMBR) to enrich for purple phototrophic bacteria (PPB) at bench (Hülsen et al., 2016). It was also in the range of traditional SBRs operated with conventional activated sludge (Mace and Mata-Alvarez, 2002). An operation at 4 cycles d^-1^ may be foreseen. Decreasing the settling phase length would lead to selectively retain the biomass fraction with higher settling property, with granulation potentialities. This is a typical approach to form a granular sludge out of flocculent activated sludge (de Kreuk and van Loosdrecht, 2006; Lochmatter and Holliger, 2014; Winkler et al., 2018).

The settling ability increased along the SBR operation, with a settled biomass fraction raising from 12% (SBR1) to 97% (SBR2-3). Amplicon sequencing revealed that the settled biomass accounted for a 3-fold higher relative abundance of PNSB (80% as sum of *Rhodobacter*, *Rhodopseudomonas*, and *Blastochloris*) than the non-settled biomass (25%) (Figure 4). Together with the phase-contrast microscopy measurements, this highlighted that PNSB are capable of forming bioaggregates with good settling properties. Such increased settling ability links to a more efficient separation of the PNSB biomass from the treated bulk liquid, thus facilitating the downstream processing to recover and valorize the PNSB biomass rich in nutrients for biorefinery purposes.

### 4.2 A high, simultaneous removal of C-N-P nutrients was achieved by the PNSB biomass

High performances of organic matter (96% COD removal at a volumetric rate of 1.1 kg COD d^-1^ m^-3^), ammonium (77% N-removal at 113 g N d^-1^ m^-3^), and orthophosphate (73% P-removal at 15 g P d^-1^ m^-3^) removal were obtained under operation with a single anaerobic reaction phase using the PNSB process. Conventionally, a sequence of anaerobic, anoxic, and aerobic conditions is needed for full BNR in activated sludge or granular sludge (Barnard and Abraham, 2006; de Kreuk et al., 2005). The main difference relies that with PNSB single organisms can remove all nutrients by assimilation into the biomass by making use of photonic energy. BNR activated sludges make use of different microbial guilds of nitrifiers, denitrifiers, polyphosphate- and glycogen-accumulating organisms among others to remove all nutrients biologically. In activated sludge or granular sludge SBRs, the different redox conditions should be alternated to this end. Hence, this PNSB SBR process is a very interesting compact alternative to conventional BNR systems, that enables an enhanced removal of all nutrients in a single reaction phase by managing one single predominant microbial guild, thus simplifying considerably the microbial resource management.

With the PNSB biomass, the SBR process becomes simpler in terms of sequencing operation by feeding, anaerobic reaction, settling, and withdrawal. In practice, a fill/draw phase can be envisioned in function of the settling properties of the PNSB biomass. This can result in a SBR system operated by alternation of fil/draw and reaction phases only. Energy-wise, aeration is not needed in a PNSB process, resulting in possible electricity savings. In the case of sunlight use, electricity savings will be substantial. The tank will have to be equipped with light filters to supply IR light and select for PNSB as predominant phototrophs in the mixed culture. In the case of ‘artificial’ supply of IR light, *e.g.*, with LEDs, the process economics will have to be balanced with the electrical power needed to provide the irradiance needed to run the process. A PNSB-based SBR process can become efficient by supplying IR light with LEDs immersed in the reactor tank or, *e.g.*, with floating carriers that can emit light under radiowaving Biofilm formation on light tubes will necessitate periodical cleaning to remediate shading, such as conventionally done for the maintenance of sensors used for process monitoring and control. The irradiance of 375 W m^-2^ applied in this bench-scale photoSBR is high versus of practical operation window. Sunny regions of Europe are typically characterized by an annual average sunlight irradiance of 150 W m^-2^ (Posten, 2009). Nonetheless, light can be provided synthetically in photobioreactors using, *e.g.*, immersed LED devices. The aim of this study was not to optimize the reactor design. Definitely, thermoeconomical analysis will have to be conducted to determine the optimum irradiance to supply.

This is analogical to the comparison of stirring performances in bench-scale reactors versus full-scale systems. There is definitely room to study PNSB processes at different illumination intensities and their impact on the system responses such as enrichment grades, biomass concentrations, aggregation levels, and nutrient removal performances. Recent studies published on purple phototrophic bacteria involved irradiances of *ca*. 50 W m^-2^ (Hülsen et al., 2016; Puyol et al., 2017a) which is about 8-times lower than the one used here at bench. However, no study has yet come with clear information on irradiance cutoffs and light patterns related to the economics of pilot and full-scale PNSB processes.

The volumetric removal rate of up to 1.1 kg COD d^-1^ m^-3^ achieved under the operation of SBR3 is comparable to the ranges of 0.8-2.5 kg COD d^-1^ m^-3^ reported for the PAnMBR (Hülsen et al., 2016), 0.2-1.4 kg COD d^-1^ m^-3^ for a continuous-flow stirred-tank reactor without separation of PNSB biomass (Alloul et al., 2019), and 1.2-3.2 kg COD d^-1^ m^-3^ for conventional BNR activated sludge processes (Tchobanoglous *et al*., 2003). It was nonetheless higher compared to aeration reactors, anaerobic ponds, and oxidation ditches (Tchobanoglous *et al*., 2003). Further enhancement of the COD loading rate and removal rate will be achieved by decreasing the SBR cycle time.

The difference in COD:N:P assimilation ratio between SBR1-2 (100:9.2:1.2 m/m/m) and SBR3 (100:6.7:0.9) resulted from the doubling of the acetate load in the influent. These COD:N:P assimilation ratios were in the range of ratios of 100:5.1-7.1:0.9-1.8 that have been characterized during growth of purple phototrophic bacteria (Hülsen *et al*., 2016a; Puyol *et al*., 2017b). As comparison basis, a COD-N-P assimilation ratio of 100:5:1 is theoretically used for activated sludge (Henze et al., 2000).

The oscillating periods of higher ISS content (>40%) in the PNSB biomass during SBR3 can underlie an interesting potential accumulation of inorganics in the biomass, *e.g.*, as intracellular polyphosphate. Similar ISS fractions of 30-40% have been widely detected in biomasses engineered for an enhanced biological phosphorus removal (EBPR) (Weissbrodt et al., 2014a). It is interesting here to mention that the model polyphosphate-accumulating organism (PAO) “*Candidatus* Accumulibacter” belongs to the microbially diverse betaproteobacterial family of *Rhodocyclaceae* (Weissbrodt et al., 2014b), thus semantically sharing the prefix “Rhodo-” with many of PNSB genera. “*Ca.* Accumulibacter” is taxonomically close to the genus *Rhodocyclus* (Hesselmann et al., 1999) which notably comprises the PNSB species *Rhodocyclus purpureus*. Although early unsuccessful testing of growth of “*Ca.* Accumulibacter” under light in the late 90’s by these authors indicating that “*Ca.* Accumulibacter” may have lost phototrophic machinery by evolution, it is worth noting that PNSB populations and “*Ca.* Accumulibacter” can share functional traits for polyphosphate accumulation. PNSB have been shown to store phosphorus to some extent (15% of cell mass) (Liang et al., 2010). Ecophysiological elucidation of PNSB populations for phosphorus removal is of high scientific and technological interests.

### 4.3 Acetate and wavelength gradients underlie microbial selection in the PNSB guild

Spectrophotometric measurements of the biomass by wavelength scans from 300 to 900 nm revealed absorbance peaks characteristics for carotenoids and bacteriochlorophylls in PNSB. Peaks at 805 and 850 nm are typical for bacteriochlorophylls detected *in vivo* from cells of *Rhodopseudomonas capsulata* (Madigan and Gest, 1979), whereas a peak around 865 nm is typical for *Rhodobacter* (Zubova et al., 2005). These peaks were detected across the whole experimental period, indicating the presence and selection of PNSB organisms in the process. Pigments are excellent biomarkers of phototrophic populations, and provide specificity to distinguish between them (Stomp et al., 2007). Wavelength scan analyses are therefore very efficient for a rapid measurement (at min level) of the selection of PNSB.

The 16S rRNA gene amplicon sequencing analysis provided insights at higher resolution on the composition of the PNSB guild and underlying selection phenomena. Amplicon sequencing revealed a consistent enrichment of PNSB after already the first 40 h of batch. The initial enrichment of *Rhodobacter* followed by selection of *Rhodopseudomonas* and then *Blastochloris* can be explained by competition phenomena across substrate and wavelength gradients between these genera inside the guild of PNSB.

Okubo & Hiraishi (2007) have reported a preferential selection of *Rhodobacter* under high acetate concentration (5-20 mmol L^-1^, *i.e.*, 320-1280 mg CODs L^-1^) due to its low affinity for acetate, while *Rhodopseudomonas* was enriched at lower concentrations (0.5 – 1 mmol L^-1^, *i.e.*, 32-64 mg CODs L^-1^). The initial 40-h batch was fully loaded with acetate across the whole reaction period, making this condition favourable to select for *Rhodobacter*. Instead, *Rhodopseudomonas* harbours a higher affinity (*i.e.*, lower affinity constant K_s_ of 0.11 mM for *Rhodopseudomonas vs* 0.23 mM for *Rhodobacter*) (Okubo and Hiraishi, 2007) for acetate, enabling this population to grow more efficiently than *Rhodobacter* under acetate-limited conditions. During SBR1 and SBR2, the carbon source became progressively depleted after 1.5 h of reaction phase, leaving other 2.5 h of starvation period at low residual acetate concentration. This provided *Rhodopseudomonas* with a competitive advantage for growth.

The competition between *Rhodobacter* and *Rhodopseudomonas* may also be governed by their growth rate and thus the SRT in the system. Populations of *Rhodobacter* have displayed a higher maximum growth rate (1.8-2.2 d^-1^ in an enrichment and 2.3-3.8 d^-1^ with isolates) about 2.6 times faster than *Rhodopseudomonas* on VFA (Alloul et al., 2019). Batch regimes primarily select on growth rate: organisms deploy their maximum growth rate across most of a batch period during which substrate concentrations are mostly not limiting (Rombouts et al., 2019). The organism with the highest growth rate that can be activated under the actual operation conditions is therefore preferentially selected. This underlay the selection for *Rhodobacter* first prior to the progressive establishment of *Rhodopseudomonas* along the progressive increase in SRT.

The genus *Blastochloris*, which appeared during SBR3, harbours bacteriochloropyll b (BChl b) instead of BChl a in *Rhodobacter* and *Rhodopseudomonas*. BChl b absorbs lower photonic energy at higher wavelengths (1020-1030 nm) (Hoogewerf et al., 2003) and can, therefore, interestingly survive at higher cell densities (here, around 1.5 g VSS L^-1^) with lower light penetration in the bulk liquid. Absorbance of the incident IR light increased across reactor operation with the development of a biofilm dominated by *Rhodopseudomonas* on the reactor wall and with a high concentration of PNSB biomass of up to 3.8 g VSS L^-1^ that accumulated in the reactor. According to the Beer-Lambert law, the accumulation of *Rhodobacter and Rhodopseudomonas* in the reactor and wall biofilm resulted in the absorbance of the higher-energy wavelengths in the 800-850 nm range of the IR light supplied, thus acting as wavelength filter. This possibly led the remaining lower-energy IR light at wavelengths above 850 nm passing further through the mixed liquor. The conjunction of the relatively high irradiance of 375 W m^-2^ and high SRT of 11 d were likely favorable for *Blastochloris* selection. Physiological characterizations of this genus is needed in order to better predict its competition with other members of the diverse PNSB guild like *Rhodobacter* and *Rhodopseudomonas* among others.

Overall, substrate gradients, light gradients, biofilm formation, and bioaggregation were identified as factors that triggered population selection and dynamics in the PNSB enrichment. Different lineages therefore act *de concert* inside the guild of PNSB, providing metabolic redundancy and process resilience in the case of regime shifts in the process.

### 4.4 PNSB mixed cultures biotechnologies: from bench toward process development

The development of a lab-scale SBR system enriched for PNSB opens the doors for a possible upscaling of the process. A high nutrient capture was coupled with the production of a PNSB-rich biomass. Such biomass can be valorized for, *e.g.*, proteins or PHAs productions (Alloul et al., 2018; Honda et al., 2006; Hülsen et al., 2018). The high settling ability of the biomass allows an easier solid-liquid separation of the latter from the treated water either in a compact external settler or directly in the SBR tank. This provides a definite downstream processing advantage over suspended biomass. It also overcomes the use of membrane filtration modules. The SBR regime resulted in the efficient aggregation of PNSB, underlying an enhanced settling ability and accumulation of biomass in the system. The SRT is an important process variable to control toward a stable bioprocess (Morgenroth and Wilderer, 1999). This becomes even more important in the perspective of harnessing the phosphorus removal capability of the PNSB biomass: cells saturated with phosphorus have to be effectively removed from the system such as conventionally performed by purge of excess sludge to maintain robust activated sludge or granular sludge processes operated for EBPR (Barnard and Abraham, 2006; Weissbrodt et al., 2013). The growth rate and affinity for the substrate are further important parameter to manage the selection of PNSB populations in either batch or continuous-flow reactor regimes, respectively. Similarly, light irradiance is a key operational variable since it constitutes the primary energy source for PNSB. Light penetration and distribution are directly linked to the reactor geometry. In surface water ecosystems, IR light photons are typically consumed over the first 30 cm depth. Following the Beer-Lambert law, the absorbance of light will substantially increase with the biomass concentration. Shallow reactor systems can be opportune. SBR regimes can easily be transferred from stirred-tank to any reactor design, like raceway systems (or also known as carrousel plants) currently under investigation for green and purple phototrophic mixed-culture processes (Alloul et al., submitted). The application of substrate gradients via SBR or plug-flow reactor configurations can foster biomass aggregation to sustain efficient S/L separation for biomass recovery on top of nutrient capture.

## 5 Conclusions

We investigated at bench the possibility to establish a mixed-culture PNSB process for nutrient capture from wastewater in a photobioreactor operated as a traditional simple and flexible stirred-tank SBR. This work led to the following three main conclusions:

1. SBR process conditions stimulated aggregation and accumulation (as high as 3.8 g VSS L^-1^) of a PNSB-enriched mixed culture in a fast-settling biomass that removed all nutrients biologically in a single reaction stage. The formation of compact aggregates facilitated S/L separation.
2. Nutrient removal was substantial by assimilation in the biomass, reaching simultaneously 96% of organic matter at 1.1 kg COD d^-1^ m^-3^, 77% of ammonium at 113 g N d^-1^ m^-3^, and 73% of orthophosphate at 15 g P d^-1^ m^-3^, *i.e.*, comparable to BNR activated sludge processes. Under non-COD-limiting conditions, the process reached the nutrient discharge limits set by the European Union.
3. The PNSB guild accounted for as high as 90% of the bacterial community of the sludge (*i.e.*, amplicon sequencing dataset), enabling a simple management of the microbial resource. A sequential selection between the genera *Rhodobacter*, *Rhodopseudomonas*, and *Blastochloris* was detected inside PNSB, allowing for functional redundancy in the microbiome. Next investigations should elucidate competition phenomena along growth rates, substrate affinities, and wavelength gradients across the mixed liquor.

For engineering practice, process analysis should cover the technological and economical aspects related to light supply in the bioreactor. Besides wastewater treatment, the value of the PNSB-based mixed-culture SBR process will reside in opportunities for water and resource recovery by valorization of the retained, concentrated, and nutrient-rich PNSB biomass.

## Supplementary material

- Supplementary material 1: Symbols and formula
- Supplementary material 2: Correlation between VSS and absorbance measurements
- Supplementary material 3: Nutrient and biomass concentrations and compositions
- Supplementary material 4: Parameter fit in Aquasim along 40-h batch and SBRs 1-3
- Supplementary material 5: V3-V4 16S rRNA gene amplicon sequencing time series

## Acknowledgements

This study was financed by the tenure-track start-up grant of the Department of Biotechnology of the Faculty of Applied Sciences of the TU Delft (David Weissbrodt, PI). Abbas Alloul was supported as doctoral candidate from the Research Foundation Flanders (FWO-Vlaanderen; strategic basic research; 1S23018N). We acknowledge the technical assistance of Udo van Dongen, Rhody Broekman, and Zita van der Krogt, successively, with the reactor infrastructure in the fermentation facility, Dirk Geerts and Rob Keerste with SCADA, as well as Ben Abbas on molecular biology.

This manuscript will be deposited as pre-print in bioRxiv.

