## Supplementary material for "Enriching and aggregating purple non-sulfur bacteria in an anaerobic sequencing-batch photobioreactor for nutrient capture from wastewater"

### Supplementary material 1:

### Symbols and Equations

Table SM1.1. Symbols used in calculation and model computation formula. Substrate concentrations were expressed in COD (organic matter), $N\text{-}NH_4^+$ (ammonium), and $P\text{-}PO_4^{3-}$ (orthophosphate) based units. Biomass concentrations were expressed in COD based units.

| Symbols | Units | Definition |
|---|---|---|
| $C_S$ | kg S $m^{-3}$ or mg S $L^{-1}$ | Concentration of substrate in the reactor (state variable) |
| $C_{S,inf}$ | kg S $m^{-3}$ or mg S $L^{-1}$ | Concentration of substrate in the influent |
| $C_{S,0}$ | kg S $m^{-3}$ or mg S $L^{-1}$ | Concentration of substrate at beginning of reaction phase |
| $C_{S,end}$ | kg S $m^{-3}$ or mg S $L^{-1}$ | Concentration of substrate at end of reaction phase |
| $C_{S,eff}$ | kg S $m^{-3}$ or mg S $L^{-1}$ | Concentration of substrate in the effluent |
| $C_X$ | kg X $m^{-3}$ or mg X $L^{-1}$ | Concentration of biomass in the reactor (state variable) |
| $C_{X,eff}$ | kg X $m^{-3}$ or mg X $L^{-1}$ | Concentration of biomass in the effluent |
| HRT | d or h | Hydraulic retention time |
| $K_S$ | kg S $m^{-3}$ or mg S $L^{-1}$ | Half-saturation affinity constant for substrate |
| $m_S$ | kg S $d^{-1}$ $kg^{-1}$ X | Biomass maintenance rate |
| $M_S$ | kg S | Mass of substrate |
| $M_X$ | kg X | Mass of biomass |
| $\mu$ or $q_X$ | kg X $d^{-1}$ $kg^{-1}$ X = $d^{-1}$ | Biomass specific growth rate |
| $N_{cycles}$ | cycles $d^{-1}$ | Number of SBR cycles per day |
| $\eta_S$ | % | Percentage of nutrient removal |
| $q_S$ | kg S $d^{-1}$ $kg^{-1}$ X | Biomass specific rate of substrate consumption |
| $Q_{inf} = Q_{eff} = Q$ | $m^3$ $cycle^{-1}$ or L $cycle^{-1}$ | Volume of influent fed and effluent withdrawn per SBR cycle |
| $Q_{sample}$ | $m^3$ $cycle^{-1}$ or mL $cycle^{-1}$ | Volume of mixed liquor samples collected during reaction phase |
| $Q_{purge}$ | $m^3$ $cycle^{-1}$ or mL $cycle^{-1}$ | Volume of mixed liquor purged at the end of reaction phase ($Q_{purge} = 0$ if SRT let freely evolve) |
| $r_S$ | kg S $d^{-1}$ $m^{-3}$ | Apparent volumetric rate of nutrient removal |
| $R_S$ | kg S $d^{-1}$ | Total rate of nutrient removal |
| SRT | d | Sludge retention time |
| t | d or h | time |
| $t_{cycle}$ | h or d | SBR cycle time length |
| V | $m^3$ or L | Volume |
| $V_{inf}$ | $m^3$ or L | Volume of influent |
| $V_r$ | $m^3$ or L | Working volume of the reactor |
| $V_{eff}$ | $m^3$ or L | Volume of effluent |
| VER | % | Volume exchange ratio |

Table SM1.2. Equations used in calculations and model computations. Definitions and units of symbols are available in Table SM2.1 above.

Hydraulic retention time in a SBR, HRT (d or h)

$$HRT = \frac{V_r}{Q_{inf}\, N_{cycles}} \quad \text{(eq. 1)}$$

Sludge retention time in a SBR, SRT (d)

$$SRT = \frac{C_x\, V_r}{(Q_{eff}\, C_{x,eff} + Q_{purge}\, C_x + Q_{sample}\, C_x)\, N_{cycles}} \quad \text{(eq. 2)}$$

Apparent volumetric rate of nutrient removal, rs (kg S $d^{-1}$ $m^{-3}$)

$$rs = \frac{(Cs,\inf Qinf - Cs,eff\ Qeff)\ Ncycles}{Vr} = \frac{(Cs,\inf - Cs,eff)\ Q\ Ncycles}{Vr} \equiv \frac{Cs,0 - Cs,end}{tcycle} \quad \text{(eq. 3)}$$

with:

$$Cs,0 = Cs,\inf\frac{Vinf}{Vr} + Cs,end\frac{Vr - Vinf}{Vr} = Cs,\inf VER - Cs,end\ (1 - VER)$$

$$Cs,inf = Cs,0\frac{1}{VER} - Cs,end\frac{1 - VER}{VER}$$

$$Cs,eff = Cs,end$$

$$VER = \frac{Vinf}{Vr} = \frac{Veff}{Vr}$$

Percentage of nutrient removal, ηs (%)

$$\eta s = \frac{Cs,inf - Cs,eff}{Cs,inf}100 = \left(1 - \frac{Cs,eff}{Cs,inf}\right)100 = \frac{Cs,0 - Cs,end}{Cs,0 - Cs,end\ (1 - VER)}100 \quad \text{(eq. 4)}$$

Total rate of nutrient removal, Rs (kg S $d^{-1}$)

$$|\ Rs\ | = \frac{dMs}{dt} = Vr\ \frac{dCs}{dt} = Vr\ rs \quad \text{(eq. 5)}$$

Substrate consumption balance during a batch phase (at constant $V_r$)

$$\frac{dC_s}{d_t} = q_s\, C_x = q_{s,max}\, \frac{C_s}{C_s + K_s}\, C_x \quad \text{(eq. 6)}$$

Biomass production balance during a batch phase (at constant $V_r$)

$$\frac{dC_x}{d_t} = \mu\, C_x = (q_s - m_s)\, Y_{sx,max}\ C_x = (q_{s,max}\, \frac{C_s}{C_s + K_s} - m_s\, )\, Y_{sx,max}\, C_x \quad \text{(eq. 7)}$$

Herbert-Pitt equation for substrate allocation for growth and maintenance

Biomass specific rates of substrate consumption, $q_s$ (kg S $d^{-1}$ $kg^{-1}$ X)
Biomass specific growth rate, μ (kg X $d^{-1}$ $kg^{-1}$ X = $d^{-1}$)
Biomass maintenance rate, $m_s$ (kg S $d^{-1}$ $kg^{-1}$ X)

$$q_s = \frac{1}{Y_{x/s}}\mu + m_s \qquad \text{or} \qquad \mu = (q_s - m_s)\, Y_{x/s} \quad \text{(eq. 8)}$$

$$\mu_{max} = \left(q_{s,max} - m_s\right) Y_{x/s} \approx q_{s,max}\, Y_{x/s} \quad \text{with } q_{s,max} \gg m_s \quad \text{(eq. 9)}$$

#### Supplementary material 2:
#### Correlation between VSS and absorbance measurements

The correlation between the absorbance measurement and the VSS concentration of the PNSB-enriched biomass is described in Figure SM1.1 hereafter. It displays the experimental data of three measurement series with the linear regression line. Such a correlation analysis should checked on regular basis across a long-term experimental period in function of the composition of the biomass. It should also be done for every new experiment, since the content of pigments and of intracellular storage compounds of the PNSB biomass can vary from case to case, based on environmental conditions tested. Absorbance measurements of biomass are mainly applicable with biomass in suspension and at relatively low concentrations. It was mainly valid here for the first 40-h batch period and SBR1. As soon as biomass aggregates, traditional gravimetry measurements via TSS, ISS and VSS as described in standard methods are more effective. This was applied on SBR2 and SBR3. One should nonetheless keep in mind that gravimetry measurements do not differentiate cells and organic intracellular polymers such as PHAs.

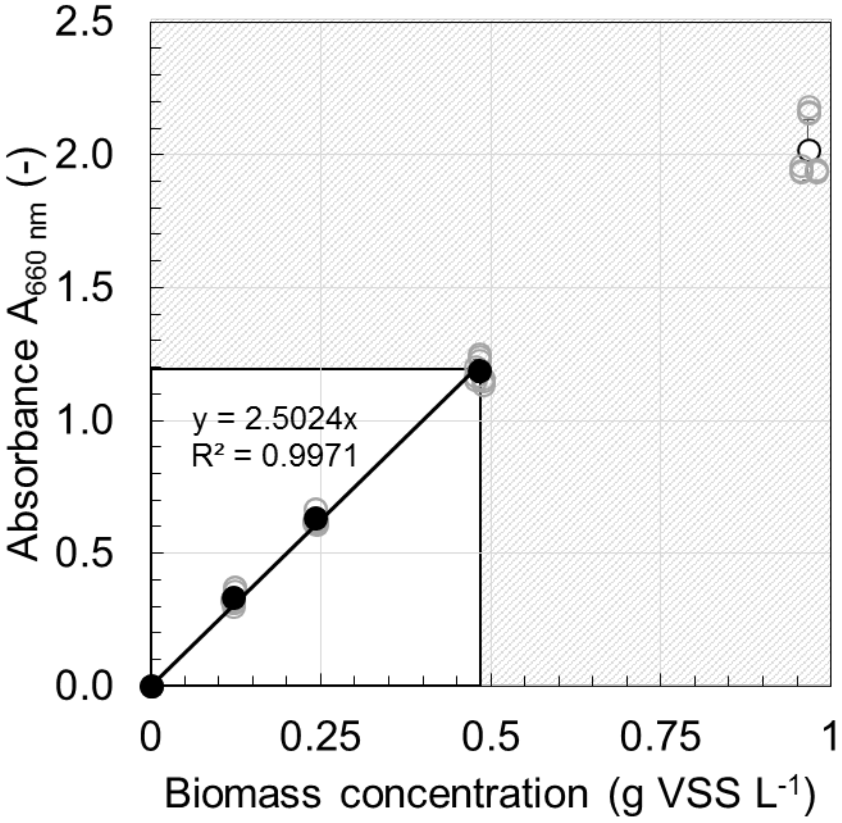


Figure SM2.1. Correlation between the absorbance at 660 nm and the VSS concentration of the PNSB-enriched biomass, based on dilution series of mixed liquor taken from the reactor (2×, 4×, 8×, 16× diluted). The linear regression of serie 1 is y = 0.3995x, serie 2 is y = 0.3856 x and serie 3 is y = 0.4124 x. For absorbance measurements, mixed-liquor samples were diluted to fit in the range from 0.2-1.2 absorbance units. On this abosrbance window the correlation to VSS was considered as linear. Absorbance data were mainly accurate measurements for the 40-h batch and SBR1 where the biomass was low concentrated and in suspension. In SBR2 and SBR3, the biomass aggregated and VSS measurements were much more accurate.

#### Supplementary material 3:

#### Dynamics of nutrient and biomass concentration and composition

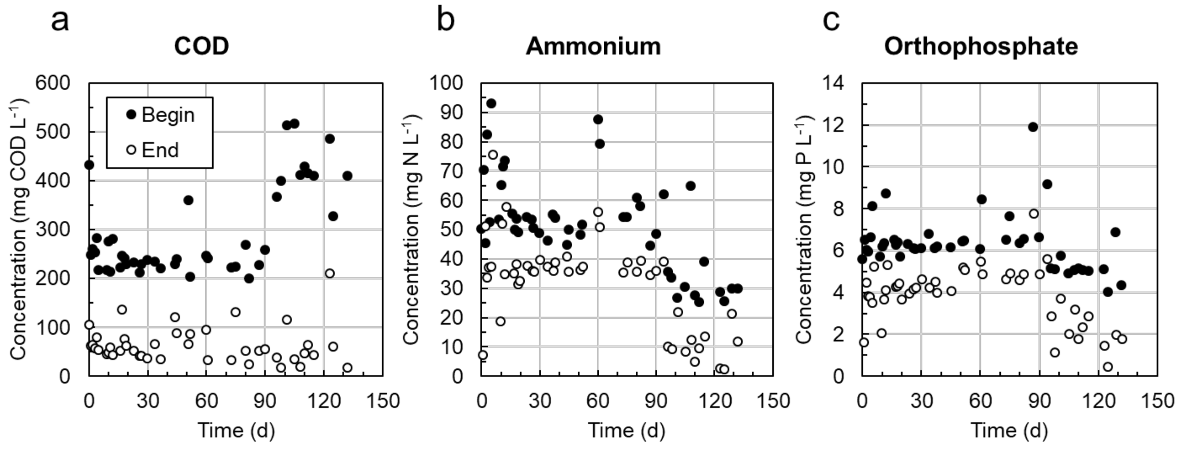


Figure SM3.1. Evolutions of COD (**A**), ammonium (**B**) and orthophosphate (**C**) removal across SBR1 (1st month), SBR2 (2nd-3rd months), SBR3 (4th-5th months). Concentrations at begin (*black dots*) and end (*white dots*) of the reaction phases of the SBRs are displayed. While acetate was fully removed, the remaining COD primarily related to EDTA present in the medium (*ca.* 50 mg COD $L^{-1}$).

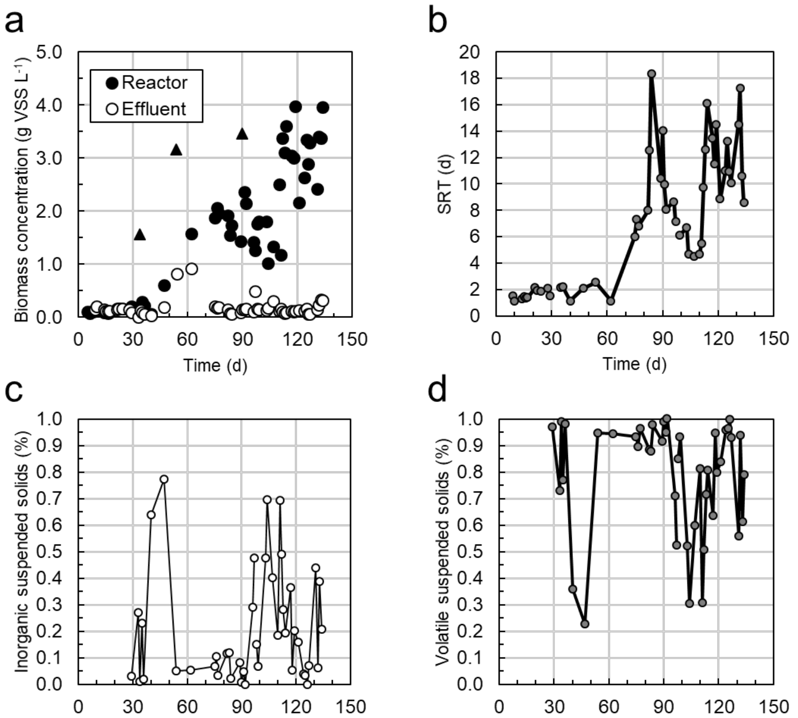


Figure SM3.2 Evolutions of (**A**) biomass concentrations in the reactor (*black dots*; *black triangles* relate to wall biofilm manual resuspension) and effluent (*white dots*); (**B**) sludge retention time (SRT); (**C-D**) fractions of inorganic (ISS) and volatile (VSS) suspended solids across SBR1 (1st month), SBR2 (2nd-3rd months), SBR3 (4th-5th months). The SRT was let freely evolve over the experimental period, without controlled purged of the mixed liquor.

### Supplementary material 4:

#### Parameter fit in Aquasim along the 40-h batch and SBRs 1-3

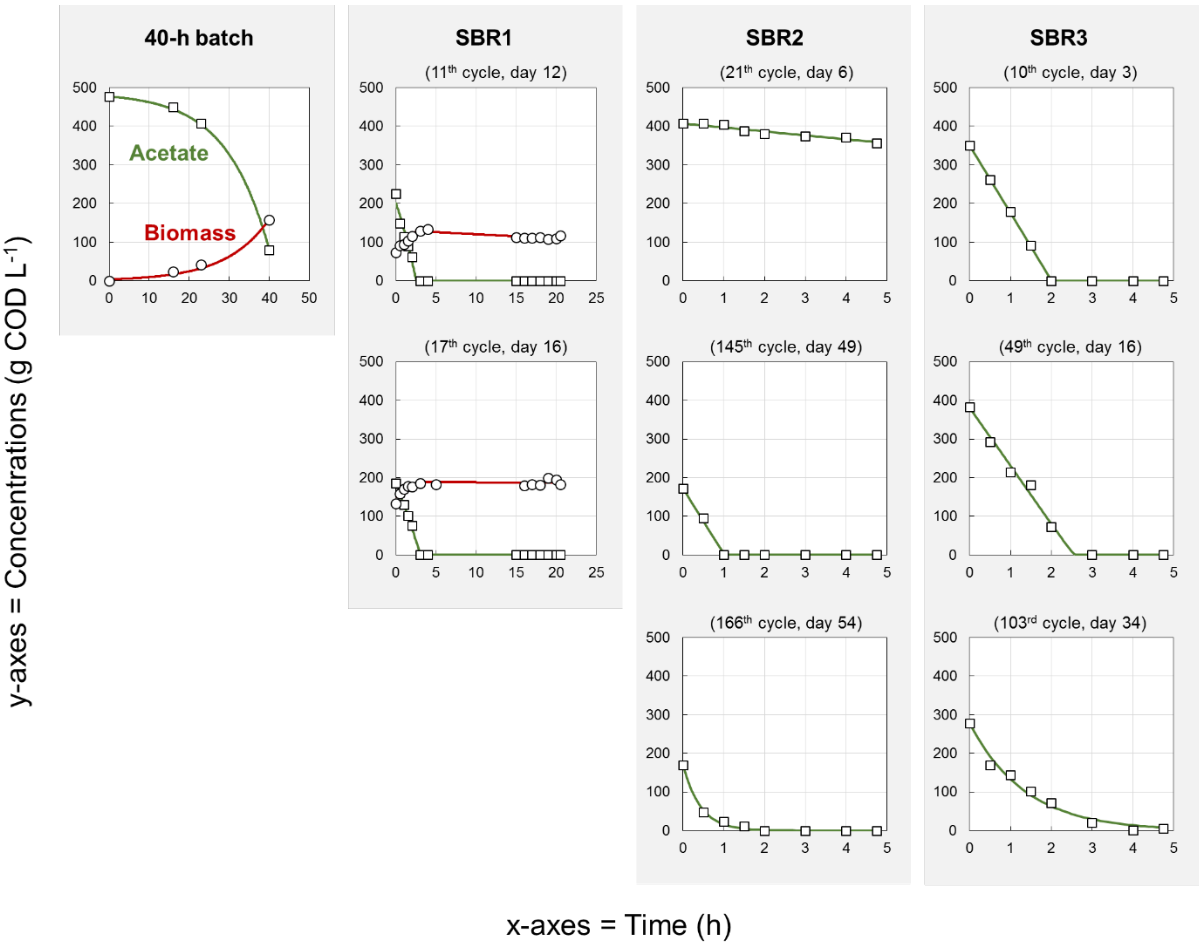


Figure SM4.1. Conversion dynamics during the 40-h batch and during the batch reaction phase of selected cycles in SBR1, SBR2, and SBR3 used for the computation of basic kinetic and stoichiometric parameters. Experimental data of biomass growth and COD depletion are displayed with the fitted Aquasim model along the batch reaction phase.

### Supplementary material 5:

### Detailed time series of V3-V4 16S rRNA gene amplicon sequencing

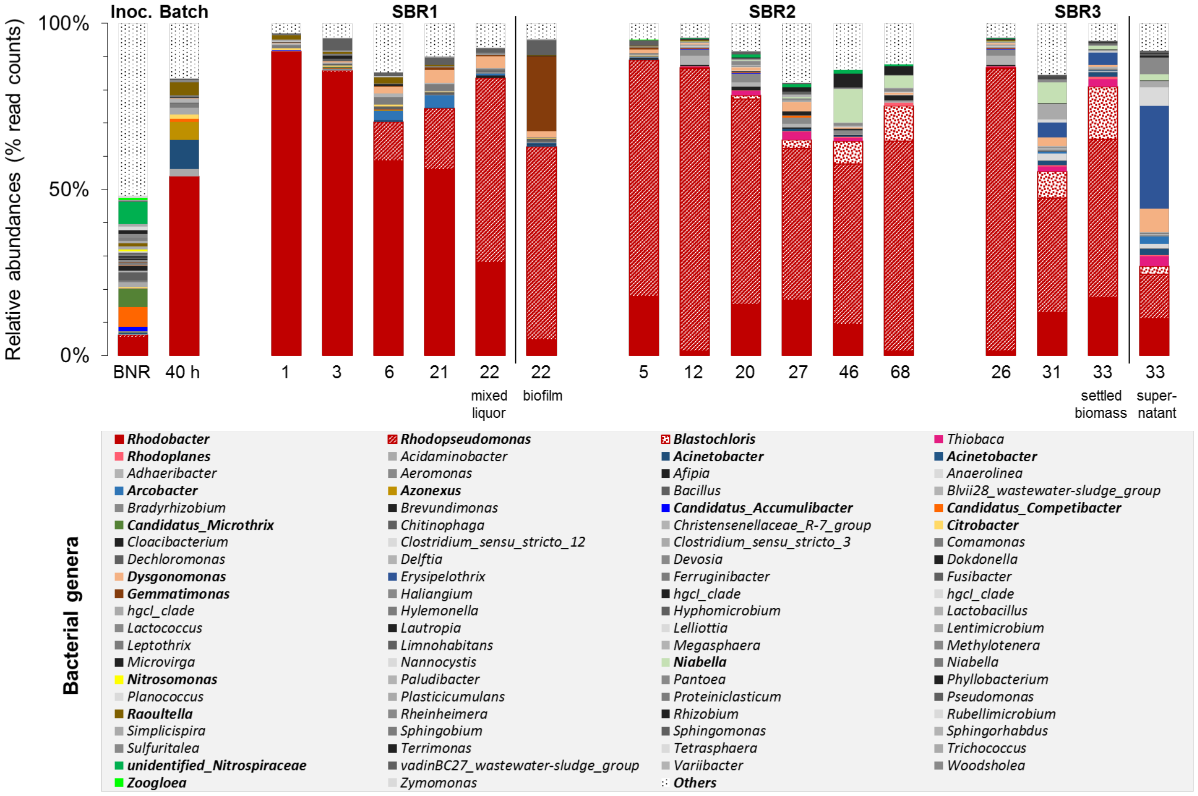


Figure SM5.1. Detailed time series of V3-V4 16S rRNA gene amplicon sequencing highlighting side populations evolving along the enrichment of PNSB in the mixed-culture process. The traditional BNR populations of the inoculum got rapidly outcompeted during the first batch after inoculation.
